# Proteolytic processing of both RXLR and EER motifs in oomycete effectors

**DOI:** 10.1101/2024.04.16.589758

**Authors:** Lin Xu, Shumei Wang, Wei Wang, Haixia Wang, Lydia Welsh, Petra C Boevink, Stephen C Whisson, Paul RJ Birch

## Abstract

Arg-any amino acid-Leu-Arg (RXLR) effectors are central oomycete virulence factors that target diverse host proteins and processes to suppress plant immunity. Relatively little is known about how they are processed post-translationally before delivery into host cells. Proteolytic cleavage at the RXLR motif was observed to occur prior to secretion in all *Phytophthora infestans* effectors tested, suggesting it is a general rule, and was observed to occur between the leucine and the second arginine. There was no cleavage of a naturally occurring second RXLR motif in a structured region of Pi21388/AvrBlb1, or one introduced at a similar position in effector Pi04314, in keeping with the motif being positionally constrained, potentially to disordered regions closely following the signal peptide. Remarkably, independent proteolytic cleavage of the Glu-Glu-Arg (EER) motif, often found immediately downstream of the RXLR, was also observed in diverse effectors, occurring immediately after the arginine. Expression of full-length effectors in host plant *Nicotiana benthamiana* revealed that, although secreted, they were poorly processed, suggesting that RXLR and EER cleavage does not occur in all eukaryotic cells. Our observations indicate that, whether possessing both RXLR and EER, or either motif alone, these effectors are likely proteolytically processed prior to secretion in all cases.

## Introduction

Diseases caused by plant pathogens and pests result in a considerable threat to food security, including up to 23% losses of the five most significant food crops (Savary et al 2019). Amongst the most economically significant disease agents are fungal and oomycete (filamentous) pathogens. The oomycete genus *Phytophthora* includes some of the most devastating plant pathogens (Derevnina et al 2016; Kamoun et al 2015). For example, *P. infestans*, causing potato and tomato late blight, precipitated the Irish potato famines of the 19^th^ century. It remains the most damaging potato and tomato disease globally (Kamoun et al 2015; Fry et al 2015).

*Phytophthora* spp. secrete ‘effector’ proteins that act either outside (apoplastic effectors) or are delivered to the inside (cytoplasmic effectors) of living plant cells. Prominent amongst cytoplasmic effectors are a class containing the conserved Arg-any amino acid-Leu-Arg (RXLR) motif (Rehmany et al 2005) located closely downstream of the signal peptide. RXLR effectors target multiple proteins and processes at diverse locations inside host cells to suppress immunity (He et al 2020; Fabro 2021; Petre et al 2021; McLellan et al 2022; Wang et al 2023a).

Many filamentous pathogens, including *P. infestans*, form haustoria, hyphal infection structures that are intimately associated with living plant cells. Haustoria are sites of cross-kingdom molecular exchange and, as such, represent key battle grounds that determine host susceptibility or resistance (Boevink et al 2020; Bozkurt and Kamoun 2020; King et al 2023). RXLR effectors have been shown to enter plant cells following their unconventional secretion from haustoria. In contrast, although also secreted from haustoria, apoplastic *P. infestans* effectors follow the canonical ER-to-Golgi pathway that is sensitive to the inhibitor brefeldin A (BFA) (Wang et al 2017; 2018). Unconventional secretion of cytoplasmic effectors and conventional secretion of apoplastic effectors has also been observed for the fungal pathogen *Magnaporthe oryzae* (Giraldo et al 2013). More recently, it has been reported that *P. infestans* RXLR effectors can be taken into plant host cells via clathrin-mediated endocytosis (CME) (Wang et al 2023b). Similarly, *M. oryzae* cytoplasmic effectors have also been observed to enter plant cells via CME (Oliveira-Garcia et al 2023), hinting at a potential universal strategy employed by haustoria-forming filamentous pathogens (Wang et al 2023c).

The RXLR motif is required for effector translocation into plant cells (Whisson et al 2007). However, its precise role has been difficult to elucidate and has often led to controversy (Ellis and Dodds 2011; Boevink et al 2020; Bozkurt and Kamoun 2020). The RXLR motif was reported to bind to phosphoinositide-3-phosphate (PI3P) on the outer surface of plant cells, promoting uptake in a pathogen-autonomous manner (Kale et al 2010). However, pathogen-independent uptake was drawn into question (Wawra et al 2013; Wang et al 2017), as was the PI3P-binding of the RXLR motif (Wawra et al 2012; Yaeno et al 2012). Indeed, the RXLR motif has been reported to be a site of proteolytic cleavage prior to effector secretion (Wawra et al 2017), implying that it is not responsible for binding to PI3P on the outer surface of the plant cell membrane.

Cleavage at the RXLR motif is reminiscent of proteolytic cleavage at an equivalent motif, RXLXE/D/Q, also called the *Plasmodium* export element (PEXEL), in effectors of the malaria parasite that are delivered into host blood cells (Boddey et al 2010; Russo et al 2010). As is apparent with the RXLR motif (Win et al 2007), the PEXEL motif is positionally constrained to within 40 amino acids after the signal peptide (SP) cleavage site. Its spatial conservation and rapid proteolytic cleavage, followed by acetylation of the N-terminus, are important for effector secretion via a specialised export pathway (Boddey et al 2016).

In addition to the RXLR motif, many *Phytophthora* cytoplasmic effectors also contain a conserved Glu-Glu-Arg (EER) motif, immediately downstream of the RXLR, which has again been implicated in effector delivery into host cells (Whisson et al 2007). Moreover, a ‘WY’ domain represents a conserved structural fold within the C-terminal half of many RXLR effectors that contributes to host target specificity (Boutemy et al 2011; Win et al 2012; Li et al 2023; Bentham et al 2023). Interestingly, a number of expressed effectors from *Bremia lactucae* that are predicted to contain the WY structural fold and are recognised by host resistance proteins only contain the EER motif (Wood et al 2020). Given that there are functionally characterised effectors containing only the RXLR motif, this raises the possibility that both motifs are effector processing sites (Wang et al 2023a).

In this work we set-out to investigate whether cleavage at the RXLR motif is widespread in *P. infestans* effectors. We discovered that cleavage is indeed evident in effectors containing RXLR-only or RXLR-EER motifs. Moreover, we observed that cleavage occurs after the leucine in the RXLR motif, rather than the second arginine as previously reported (Wawra et al 2017). We confirm that cleavage of the RXLR is constrained by its location after the SP. Unexpectedly, we discovered that the EER motif is also cleaved after the arginine, revealing that both RXLR and EER are sites for proteolytic processing.

## Materials and Methods

### *Phytophthora infestans* culture and *Nicotiana benthamiana* inoculation

*A. P. infestans* wild type (WT) strain 2006-3928A (hereafter called 3928A) was obtained from the James Hutton Institute *Phytophthora* culture collection (https://ics.hutton.ac.uk/hutton-collections/#/pathogens). The WT strain was cultured on rye agar medium (Tumwine *et al*., 2000) containing ampicillin (Amp, 50 μg/mL) and pimaricin (Pim, 20 μg/mL), and transgenic lines were cultured on rye agar medium containing Amp, Pim and geneticin (G418, 10 μg/mL) at 19 °C in the dark. To prepare inoculum, 10-14 day-old *P. infestans* culture plates were flooded with sterile distilled water (dH_2_O) to wash off sporangia from the sporangiophores. The spore suspension was filtered through 100 μm nylon mesh sterile cell strainers (Fisher Scientific) into 50 mL sterile centrifuge tubes (Sarstedt) and centrifuged at 1600 × g for 10 min. The number of sporangia was counted using a hemocytometer and the concentration of the inoculum for *N. benthamiana* leaf infection was adjusted by dilution in dH_2_O to yield 8 × 10^4^ sporangia mL^-1^. Detached and wounded *N. benthamiana* leaves were inoculated with 10 μL droplets of the sporangia suspension. Inoculated leaves were incubated on moist tissue in sealed clear plastic boxes at 19 °C in the dark for the first 24 h, then under a normal day-night light regime (Kunjeti *et al*., 2016; He *et al*., 2018).

### Vector construction for *P. infestans* transformation

The vector used to construct plasmids for co-expressing monomeric red fluorescent protein (mRFP) tagged effectors and cytoplasmic green fluorescent protein (GFP) in *P. infestans* was pPL-RAG (Wang et al 2017). Coding sequences (without stop codons) of Pi09216 (XP_002903512), Pi04314 (XP_002905959), Pi21388/IPIO1/AvrBlb1 (XP_002895051), Pi22926 (XP_002901758), Pi09160 (XP_002903475), Pi04145 (XP_002905847) were amplified from the genomic DNA of *P. infestans* WT 3928A using gene-specific primers that contain the NotI restriction enzyme recognition sequence (Supplementary Table S1). The attempt to amplify the sequence of Pi04097 (XP_002905808) directly using gene specific primers failed due to sequence similarities among members of the Avrblb2 family, to which Pi04097 belongs, so a nested PCR approach was used. Primers flanking the coding sequence (Supplementary Table S1) were used in the first round PCR to amplify a longer sequence that contained Pi04097 from the genomic DNA of 3928A. This was used as template in the second round PCR to amplify the coding sequence (without stop codon) of Pi04097 using gene-specific primers that contain the NotI restriction enzyme recognition sequence (Supplementary Table S1). All the PCR products were purified (MinElute PCR purification Kit, QIAGEN) and digested with NotI (New England BioLabs, NEB). The digested PCR products were purified and ligated into NotI-digested and shrimp alkaline phosphatase (NEB) dephosphorylated pPL-RAG vector using T4 DNA ligase (NEB). Ligation products were electroporated into *Escherichia coli* DH10B Electromax electrocompetent cells (Invitrogen). Transformed cell suspensions were plated on Luria-Bertani (LB) agar medium containing 50 μg/mL Amp. Colonies were screened by colony PCR using gene-specific forward primers and a reverse primer located near the 5’ end of the mRFP gene (Supplementary Table S1). Plasmids purified from positive colonies were verified by sequencing using the mRFP reverse primer and named as pPL-RAG-effector number.

The RXLR or EER mutant vectors were generated using the QuikChange II XL site-directed mutagenesis kit (Agilent). Gene-specific mutagenic primers were designed using the QuikChange primer design program (https://www.agilent.com/store/primerDesignProgram.jsp) (Supplementary Table S1). pPL-RAG-09216, pPL-RAG-04097, pPL-RAG-04314, pPL-RAG-21388 and pPL-RAG-22926 were used as templates to yield the pPL-RAG-09216AAAA, pPL-RAG-04097AAAA, pPL-RAG-04314AAAA-EER, pPL-RAG-04314RXLR-AAA, pPL-RAG-04314RXLR-EER-RXLR, pPL-RAG-21388AAAA-EER, pPL-RAG-21388RXLR-AAA, pPL-RAG-22926AAAA-EER and pPL-RAG-22926RXLR-AAA vectors, respectively. Then pPL-RAG-04314AAAA-EER, pPL-RAG-21388AAAA-EER and pPL-RAG-22926AAAA-EER vectors were used as templates to yield the pPL-RAG-04314AAAA-AAA, pPL-RAG-04314AAAA-EER-RXLR, pPL-RAG-21388AAAA-AAA and pPL-RAG-22926AAAA-AAA vectors, respectively.

### *P. infestans* transformation

Transformation of *P. infestans* was carried out according to a modified polyethylene glycol (PEG)-CaCl_2_-lipofectin mediated method (Judelson *et al*., 1991; Avrova *et al*., 2008). A version of the protocol can be found at https://oomyceteworld.net/TransformationMethods.html. The WT used for transformation was a line regenerated from 3928A protoplasts (3928A mock) that had been through the transformation procedure but without any transforming DNA (mock transformed). The plasmid DNA used was prepared using GenElute™ HP plasmid maxiprep kit (Sigma-Aldrich). The protoplasting solution contained 5 mg/mL Extralyse (Laffort) and 4 mg/mL cellulase from *Trichoderma viride* (Duchefa Biochemie). For each transformation, at least 50 μg plasmid DNA in 57.1 μL sterile dH_2_O was mixed with 85.7 μL Lipofectin (Invitrogen). Protoplasts were regenerated in LBSM (lima bean-sucrose-mannitol) liquid medium at 19 °C in the dark for 48 h, collected by centrifugation at 1000 × g for 5 min, then spread onto rye agar medium containing 2.5 μg/mL G418 for selection. G418-resistant colonies typically appeared after 8 to 14 days, and they were transferred to and maintained on rye agar medium containing 10 μg/mL G418.

### Vector construction for expression *in planta*

Pi09216_1-175_, Pi04314_1-154_, Pi21388_1-152_ and Pi22926_1-198_ (full length coding sequences without stop codons), Pi09216_38-175_, Pi04314_57-154_, Pi21388_55-152_ and Pi22926_47-198_ (coding sequences immediately after the RXLR motif to the end, without stop codons), and Pi04314_68-154_, Pi21388_73-152_ and Pi22926_53-198_ (coding sequences from immediately after the EER motif, without stop codons) were amplified from pPL-RAG-09216, pPL-RAG-04314, and pPL-RAG-21388 vectors, respectively. Gene-specific primers contained flanking Gateway (Invitrogen) recombination sequences (Supplementary Table S1). PCR products were purified and recombined into the pDONR201 entry vector (Invitrogen) using Gateway cloning (BP reaction) to generate donor vectors. These donor vectors were recombined with destination vectors pK7RWG2 (C-mRFP) by Gateway cloning (LR reaction) to generate overexpression vectors. Sequence-verified plasmids were electroporated into *Agrobacterium tumefaciens* strain GV3101 for transiently overexpressing mRFP fusion proteins in *N. benthamiana*.

### *Agrobacterium*-mediated transient expression

*A. tumefaciens* containing overexpression vectors were grown at 28 °C overnight with shaking at 170 rpm, in yeast extract and beef (YEB) liquid medium with appropriate antibiotics. The bacteria were pelleted by centrifuging at 3000 × g for 10 min and then resuspended in infiltration buffer (10 mM 2-(*N*-morpholino) ethanesulfonic acid (MES), 10 mM MgCl_2_). The concentration was adjusted as required to the appropriate optical density at 600 nm (OD600). The OD600 used for confocal imaging was 0.01−0.05, and for immunoblots was 0.5. The required volume of bacteria suspension was supplemented with 200 mM acetosyringone and incubated at room temperature in the dark for at least 2 h before *N. benthamiana* leaf infiltration.

### Protein extraction and immunoprecipitation

To extract effector-mRFP fusion proteins from *P. infestans* mycelia (M) and culture filtrate (CF), sporangia of *P. infestans* transformants were collected from 10 to 14-day-old plates and cultured in 5 mL amended lima bean (ALB) liquid medium (Bruck *et al*., 1980) containing 10 μg/mL G418 at 19 °C in the dark. After 3 days, the M were transferred to 1 mL ALB liquid medium containing 10 μg/mL G418 and cultured at 19 °C in the dark. After 1 day, M were separated from the growth medium (referred to as CF) and semi-dried using tissue. The semi-dried M were thoroughly mixed with 50 μL 2 × sodium dodecyl sulfate (SDS) sample loading buffer (100 mM Tris pH=6.8, 4% SDS, 20% glycerol, 200 mM dithiothreitol (DTT), 0.2% bromophenol blue) and heated at 95 °C for 10 min. The heated sample was centrifuged at 13000 × g for 10 min and the supernatant was used for SDS-PAGE gel loading. The CF was centrifuged at 1000 × g for 5 min to remove cellular debris, and the total protein present in the supernatant was precipitated using chloroform/methanol precipitation (Fic *et al*., 2010). Briefly, four volumes of methanol, one of chloroform and four of sterile dH_2_O were added to one volume of centrifuged CF, and the mixture was vortexed and centrifuged at 4500 × g for 15 min. A white interphase formed between the methanol and chloroform liquid layers after the centrifugation. The liquid layers were discarded, two volumes of methanol were added to the remaining interphase, and the mixture was vortexed and centrifuged at 4500 × g for 10 min. The supernatant was discarded, the pellet was air dried, resuspended in 50 μL 2 × SDS sample loading buffer, and heated at 95 °C for 10 min. The heated sample was centrifuged at 13000 × g for 10 min and the supernatant was used for SDS-polyacrylamide gel (PAGE) loading.

To extract effector mRFP fusion proteins from *N. benthamiana* leaves infiltrated with *Agrobacterium* containing overexpression vectors, four leaf discs (1 cm in diameter) were snap frozen in liquid nitrogen and powdered using a TissueLyser (QIAGEN). Total proteins were extracted from the powder using 400 μL GTEN buffer (10% glycerol, 25 mM Tris-HCl pH=7.5, 1 mM ethylenediaminetetraacetic acid (EDTA), 150 mM NaCl) with 10 mM DTT, protease inhibitor cocktail (PIC), 1 mM phenylmethylsulfonyl fluoride (PMSF) and 0.2% Nonidet P-40 added. 50 μL of the extraction was mixed with 50 μL 2× SDS sample loading buffer, heated at 95 °C for 10 min and used for SDS-PAGE gel loading.

To immunoprecipitate effector mRFP fusion proteins from *N. benthamiana* leaves infected by *P. infestans* transformants, 2 g of infected leaves were ground into powder in liquid nitrogen. Total proteins were extracted as described above and the extraction was mixed with 30 μL RFP-Trap® magnetic agarose beads (Chromotek) and rotated at 4 °C overnight. The supernatant was discarded, and beads were washed five times using GTEN buffer with PIC and 1 mM PMSF added. The washed beads were resuspended in 50 μL 2× SDS sample loading buffer and heated at 95 °C for 10 min. The supernatant was magnetically separated from the beads and used for SDS-PAGE gel loading.

To immunoprecipitate effector mRFP fusion proteins from *P. infestans* CF, sporangia of *P. infestans* transformants were cultured in 10 mL ALB liquid medium containing 10 μg/mL G418 at 19 °C in the dark for 3 days. M were removed and the CF was centrifuged at 13000 × g for 10 min. Total proteins were precipitated from the supernatant using acetone precipitation (Fic *et al*., 2010). Briefly, four volumes of chilled acetone were added to one volume of the centrifuged CF and the mixture was vortexed and incubated at −20 °C overnight. The mixture was centrifuged at 4500 × g for 10 min at 4 °C, the supernatant was discarded, and the pellet was air dried. The dried pellet was dissolved in 1 mL GTEN buffer with 10 mM DTT, PIC, 1 mM PMSF and 0.2% Nonidet P-40 added, and centrifuged at 13000 × g for 10 min at 4 °C. The supernatant was mixed with 40 μL RFP-Trap® magnetic agarose beads and rotated at 4 °C overnight. The subsequent steps were the same as described above.

### SDS-PAGE and Western blot

Samples were loaded onto 12% self-cast gels; gel electrophoresis, membrane blocking and washing steps were performed as described by McLellan *et al*. (2013). Amersham^TM^ Protran^TM^ 0.45 μm NC nitrocellulose blotting membrane was used. Primary RFP antibody (α RFP, Chromotek) and histone H3 antibody (α H3, Abcam) were used at 1 : 5000 dilutions. Secondary anti-rat immunoglobulin G (IgG) horseradish peroxidase (HRP) antibody (Abcam) and anti-rabbit IgG HRP antibody (Sigma-Aldrich) were used at 1 : 5000 dilutions. Protein bands on membranes were visualized using high performance chemiluminescence film (Amersham^TM^ Hyperfilm^TM^ ECL) and Pierce™ or SuperSignal™ ECL western blotting substrate (ThermoFisher Scientific).

### Confocal microscopy

*N. benthamiana* leaves infected by *P. infestans* (4−6 dpi) or infiltrated with *Agrobacterium* containing overexpression vectors (2 dpi) were infiltrated with dH_2_O and then mounted on slides. A Nikon A1R confocal microscope was used for imaging. RFP was imaged with 561 nm excitation and emissions were collected between 570−620 nm. GFP and chloroplast auto-fluorescence were imaged with 488 nm excitation and emissions were collected between 500−530 nm and 663−738 nm respectively. The pinhole was set to 1.2 Airy units for the longest wavelength fluorophore of any combination. All images were processed and analysed using Fiji software (Schindelin *et al*., 2012).

### Liquid chromatography-tandem mass spectrometry (LC-MS/MS)

Immunoprecipitated samples from the CF of *P. infestans* transformants were analysed by SDS-PAGE. Protein bands were visualized by InstantBlue Coomassie Protein Stain (Abcam) staining. Bands of interest were excised, destained, and digested overnight (16 h) with trypsin, chymotrypsin or lys C (MS Grade, Pierce, ThermoFisher Scientific). Peptides generated by the digestion were extracted from the gel and dried in a SpeedVac (ThermoFisher Scientific). Dried peptides were resuspended with 40 μL 1% formic acid, centrifuged and transferred to HPLC vials for analysis using Ultimate^TM^ 3000 RSLCnano system (ThermoFisher Scientific) coupled to a Q Exactive^TM^ Plus Mass Spectrometer (ThermoFisher Scientific). Mass spectrometry analysis was carried out at the ‘FingerPrints’ Proteomics Facility, School of Life Sciences, University of Dundee. 15 μL samples were injected, peptides were initially trapped on an Acclaim^TM^ PepMap^TM^ 100 (C18, 100 µM × 2 cm) and then separated on an Easy-Spray^TM^ PepMap RSLC C18 column (75 µM × 50 cm) (ThermoFisher Scientific) and transferred to the mass spectrometer via an Easy-Spray source with temperature set at 50 °C. In data-dependent acquisition mode, full MS1 scans were performed at 70000 resolution followed by 15 sequential dependent MS2 scans where the top 15 ions were selected for collision-induced dissociation (CID, normalized collision energy NCE = 35.0) and analysis in the Ion Trap with an MSn AGC target of 5000. An isolation window of 2.0 m/z units around the precursor was used and selected ions were then dynamically excluded from further analysis. Mascot data software (version 2.6) was used for data analysis.

## Results

### Cleavage of the RXLR motif in RXLR-only effectors

Cleavage at the RXLR motif of PiAVR3a detected in the culture filtrate (CF) of axenically grown *P. infestans* was reported previously (Wawra et al 2017). To investigate whether such cleavage was widespread we selected two effectors that contain the RXLR motif but lack the EER motif, Pi09216 and Pi04097, each of which enhances *P. infestans* colonisation when transiently expressed in *N. benthamiana* (Wang et al 2019). Pi04097 is an AvrBlb2 family member and is recognised by the potato NLR RpiBlb2 (Oh et al 2009). Constructs expressing either the WT versions of each effector, or versions in which the RXLR was mutated to alanines (AAAA), each fused to mRFP at the C-terminus, were transformed into *P. infestans* (**Fig. 1a**). Transformants expressing the mutant forms, Pi09216AAAA or Pi04097AAAA, generated proteins of a size consistent with cleavage after the signal peptide (SP). These effector forms were detected in both the CF and mycelium samples; the absence of histone H3 in the former confirmed that the mutant effector forms were actively secreted rather than released by cell lysis (**Fig. 1b, 1c**). Transformants expressing the WT versions of Pi09216-mRFP and Pi04097-mRFP generated smaller fusion proteins consistent with cleavage at the RXLR motif (**Fig. 1b, 1c**). The smaller fusion proteins were detected in mycelium samples, suggesting that processing of the effectors occurred prior to secretion. These smaller fusion proteins were also evident in transformants expressing mutant Pi09216AAAA-mRFP samples, perhaps indicating that cleavage was reduced rather than fully prevented by the mutation. In the case of WT Pi04097-mRFP, a faint fusion protein of a size predicted to be the SP cleavage product was detected only in the mycelium sample (**Fig. 1c**). Both the WT and mutant forms of Pi09216-mRFP and Pi04097-mRFP were evident around the outside of GFP-labelled haustoria in fluorescence intensity plots, consistent with them being secreted from haustoria during infection, indicating that mutation of the RXLR motif does not prevent this (**Suppl Fig. S1a, S1b**). To test whether similar processing of the WT effectors occurred during interactions with host cells, transformants expressing the WT and mutant mRFP fusions of Pi09216 were inoculated onto *N. benthamiana* leaves and the WT and mutant Pi09216-mRFP fusion proteins were immunoprecipitated. The WT and mutant fusion proteins were similar sizes during infection and growth in culture medium (**Suppl Fig. S2a**), indicating that similar processing occurs in each case.

**Figure 1.**
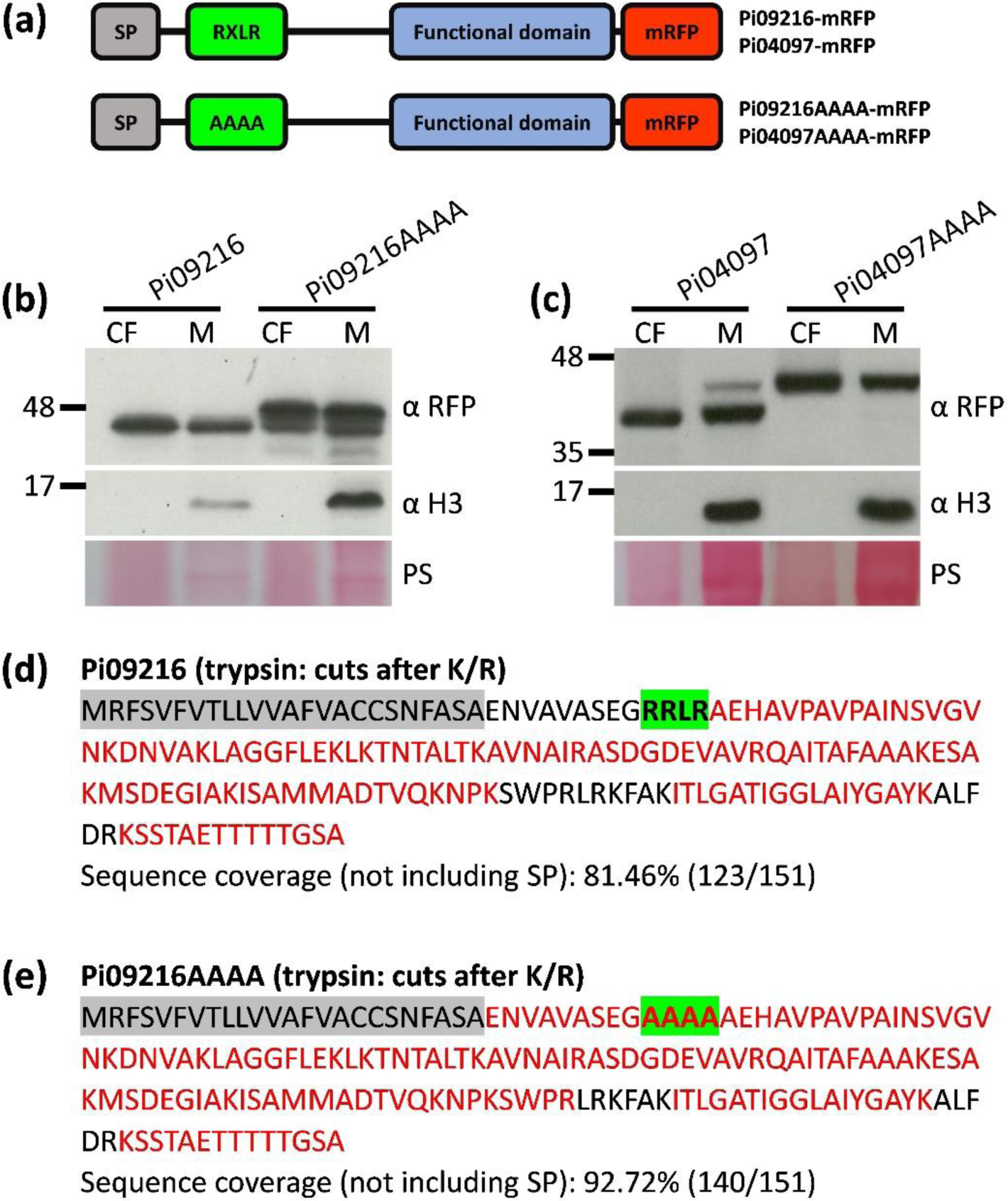
Cleavage of the RXLR motif in RXLR-only effectors. (a) Schematic representation of Pi09216 and Pi04097 wild type (WT) and RXLR mutant (AAAA) monomeric red fluorescent protein (mRFP) fusions. SP, signal peptide. Pi09216 (b) and Pi04097 (c) WT and AAAA mRFP fusions were detected in the culture filtrate (CF) and mycelium (M) using RFP antibody (αRFP). *Phytophthora infestans* histone H3 was detected using histone antibody (α H3) only in the M which indicates that the CF was not detectably contaminated by cellular material. Markers (kDa) are indicated on the left and protein loading is indicated by Ponceau stain (PS). Distribution of the LC-MS/MS identified peptides within the sequences of Pi09216 WT (d) and AAAA (e). Peptides were generated by trypsin in-gel digestion. Identified peptides are marked with red, predicted SP is shaded with grey, and the RXLR motif and its alanine replacement (AAAA) are shaded with green. K: lysine, R: arginine.

WT and mutant forms of Pi09216-mRFP were immunoprecipitated from CF following growth of *P. infestans* in axenic culture and LC MS/MS was conducted from replicate experimental samples. Whereas peptides were only detected representing sequences from the RXLR motif to the C-terminus of the WT Pi09216 (**Fig. 1d; Suppl Table S2a**), consistent with cleavage at the motif, peptides from the predicted SP cleavage site, spanning the AAAA region of mutated Pi09216, were reproducibly detected (**Fig. 1e; Suppl Table S2b**), confirming that uncleaved protein was secreted by the transformant expressing the mutated effector.

### Cleavage of effectors with both RXLR and EER motifs

To study in detail the potential for proteolytic processing of effectors containing both the RXLR and EER motifs, two well characterised effectors were selected: Pi04314, which is secreted from haustoria (Wang et al 2017) and interacts with host PP1c isoforms in the plant nucleus (Boevink et al 2016; Bentham et al 2023); and Pi21388 (IPIO1/Avrblb1) which is recognised by the potato resistance RpiBlb1 (Vleeshouwers et al 2008). Initially, WT versions of each effector, and versions in which both RXLR and EER motifs were mutated (AAAA-AAA), were expressed in *P. infestans* as mRFP fusions (**Fig. 2a**). The AAAA-AAA mutants generated fusion proteins of a size expected following cleavage at the SP. These were evident in the mycelium samples and were secreted into CF (**Fig. 2b, 2c)**. By contrast, smaller fusion proteins were observed for both WT Pi04314-mRFP and Pi21388-mRFP (**Fig. 2b, 2c).** This pattern was also evident for WT and AAAA-AAA mutant versions of RXLR-EER effector dPi22926 (**Suppl Fig. S3)**, which has been shown previously to be secreted at haustoria and translocated into host cells (Wang et al 2018). Fluorescence intensity plots indicated that both the WT and AAAA-AAA mutant forms of Pi04314, Pi21388 and Pi22926 were secreted from haustoria during infection, indicating that mutation of the RXLR and EER motifs does not prevent this (**Suppl Fig. S4**). To test whether similar processing of the WT effectors occurred during interactions with host cells, transformants expressing the WT and mutant mRFP fusions of Pi04314 were inoculated onto *N. benthamiana* leaves and the Pi04314-mRFP fusion proteins were immunoprecipitated. The WT and mutant fusion proteins were similar sizes during infection and growth in culture medium (**Suppl Fig. S2b**), indicating that similar processing likely occurs in each case.

**Figure 2.**
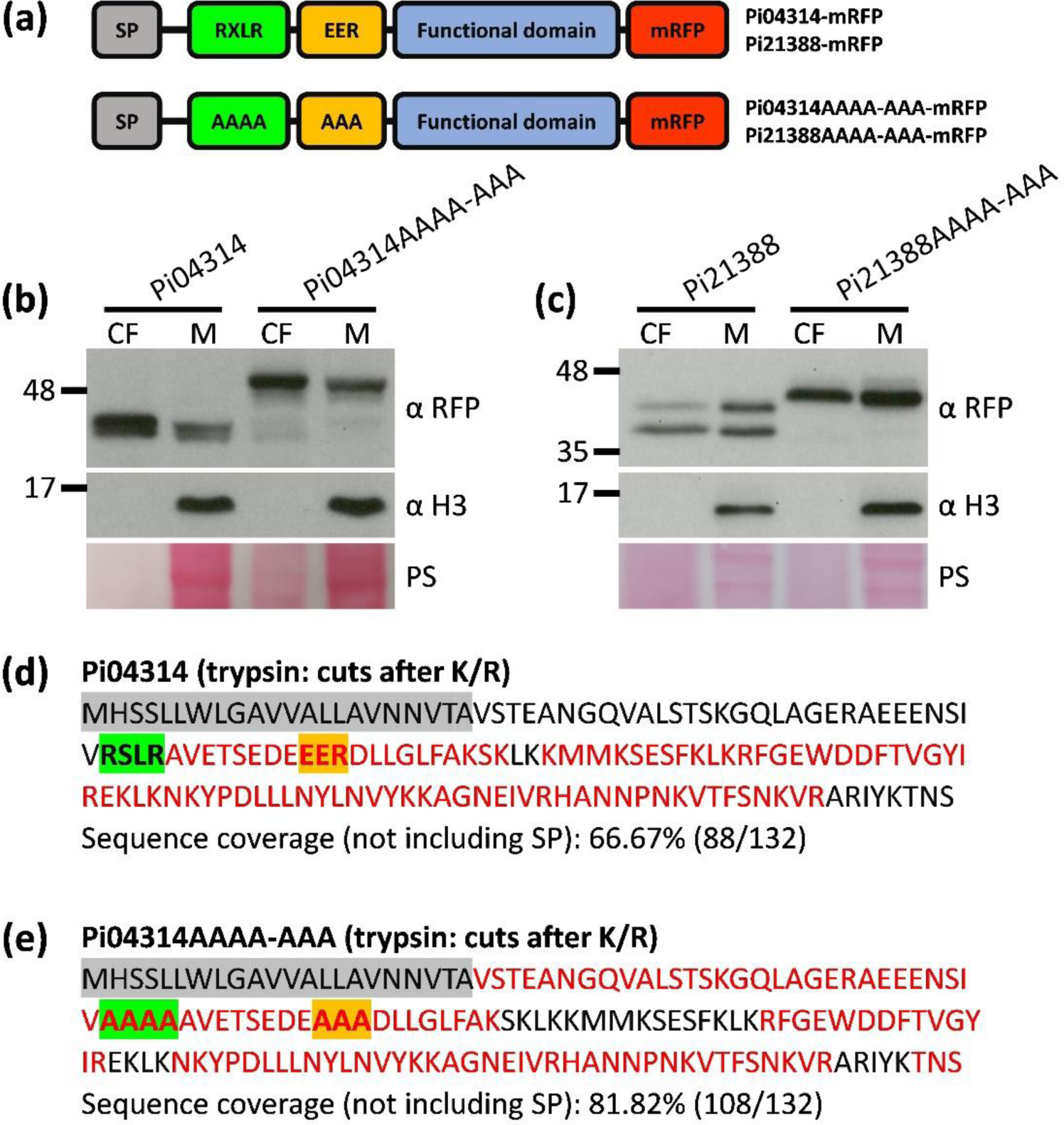
Cleavage of the RXLR motif in RXLR-EER effectors. (a) Schematic representation of Pi04314 and Pi21388 WT and RXLR-EER mutant (AAAA-AAA) mRFP fusions. Pi04314 (b) and Pi21388 (c) WT and AAAA-AAA mRFP fusions were detected in the culture filtrate (CF) and mycelium (M) using αRFP. *P. infestans* histone H3 was detected using α H3 only in the M which indicates that the CF was not detectably contaminated by cellular material. Markers (kDa) are indicated on the left and protein loading is indicated by Ponceau stain (PS). Distribution of the LC-MS/MS identified peptides within the sequences of Pi04314 WT (d) and AAAA-AAA (e). Peptides were generated by trypsin in-gel digestion. Identified peptides are marked with red, predicted SP is shaded with grey, the RXLR motif and its alanine replacement (AAAA) are shaded with green, and the EER motif and its alanine replacement (AAA) are shaded with gold.

WT and mutant Pi04314-mRFP fusion proteins were immunoprecipitated from CF following growth of *P. infestans* in axenic culture and LC-MS/MS was conducted from replicate experimental samples. Peptides were only detected from after the RXLR motif to the C-terminus of the WT Pi04314 (**Fig. 2d; Suppl Table S2c**), consistent with cleavage at the motif. By contrast, peptides spanning the AAAA and AAA regions of mutated Pi04314 were reproducibly detected (**Fig. 2e; Suppl Table S2d**), consistent with a much-reduced cleavage in the AAAA-AAA mutant effector.

### Evidence of cleavage at both RXLR and EER motifs

We noted that WT Pi21388-mRFP transformants revealed two protein bands in both mycelium and CF samples (**Fig. 2c**). Close inspection revealed that two fusion proteins were apparent also for WT Pi04314-mRFP (**Fig. 2b**) but these were closer in size. The differences in size between the fusion protein pairs produced by the transformants expressing Pi21388-mRFP and Pi04314-mRFP may reflect distance between the RXLR and EER motifs; 15 amino acids in the case of Pi21388 and 8 amino acids in the case of Pi04314-mRFP (**Fig. 3**). Two fusion protein bands were not initially evident in the case of WT Pi22926-mRFP (**Suppl Fig. S3)**. This may reflect that only 3 amino acids separate RXLR and EER motifs in Pi22926-mRFP (**Suppl Fig. S5**). To test the hypothesis that each motif was a cleavage site we mutated either the RXLR (AAAA-EER) or the EER (RXLR-AAA) motif independently for each effector (**Fig. 3a)**. As observed previously (**Fig. 2; Suppl Fig. S3)**, the double mutants (AAAA-AAA) of Pi04314-mRFP, Pi21388-mRFP and Pi22926-mRFP generated protein bands of a size consistent with cleavage at the SP in both mycelium and CF samples. Moreover, WT fusion proteins for each effector revealed smaller bands consistent with cleavage at the conserved motifs (**Fig. 3b, c; Suppl Fig. S5).** Effector forms in which the RXLR (AAAA-EER) was mutated generated two fusion products; one with predicted cleavage at the SP and one consistent with cleavage at the EER motif (**Fig. 3b, c; Suppl Fig. S5).** Effector forms in which the EER (RXLR-AAA) motif was mutated generated a dominant fusion product of a size predicted following cleavage at the RXLR motif (**Fig. 3b, c; Suppl Fig. S5).** Each of the single-motif mutant transformants expressed mRFP fusion proteins that were secreted at haustoria during infection (**Suppl Fig. S6).**

**Figure 3.**
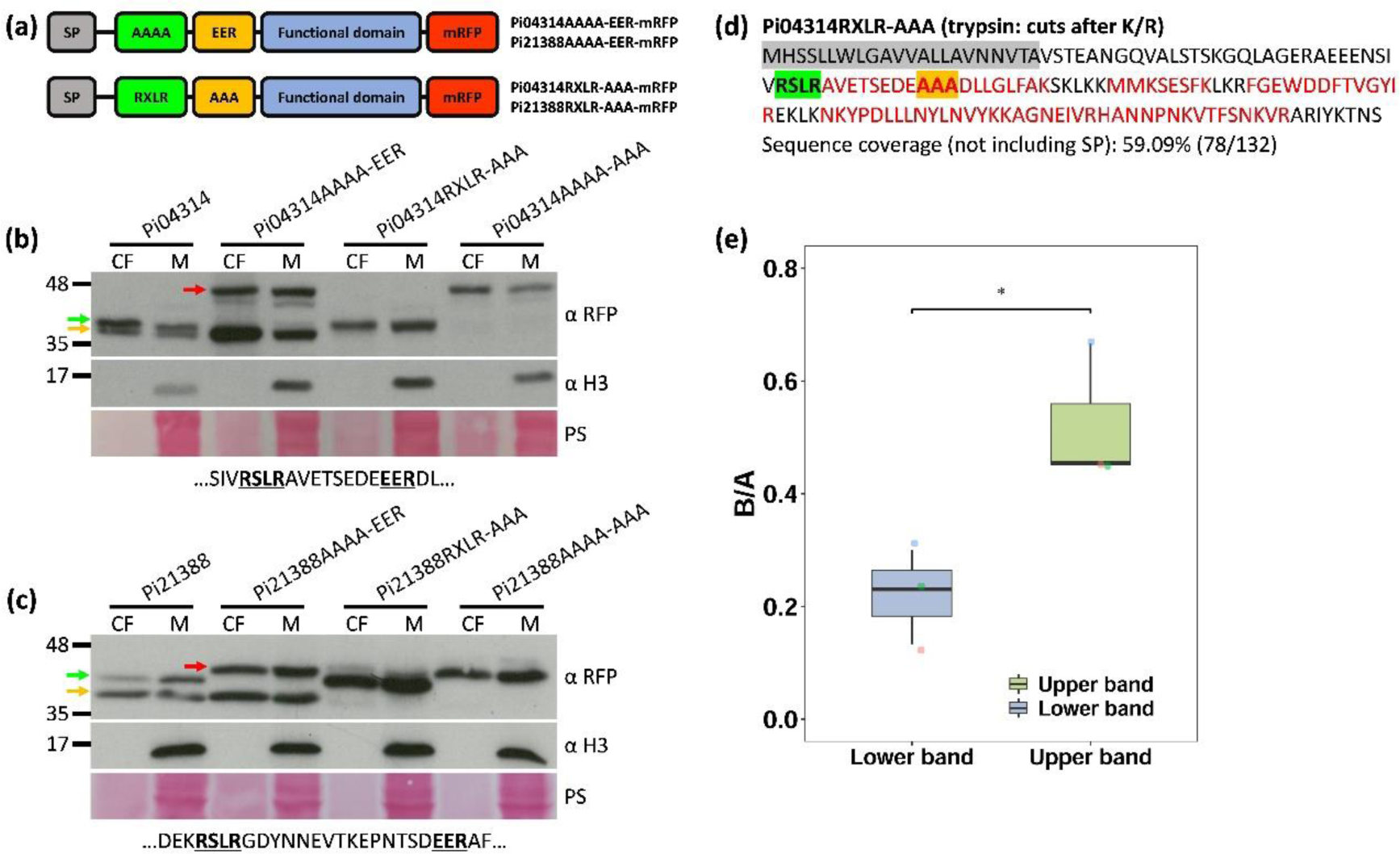
Evidence of cleavage at both RXLR and EER motifs. (a) Schematic representation of Pi04314 and Pi21388 RXLR mutant (AAAA-EER) and EER mutant (RXLR-AAA) mRFP fusions. Pi04314 (b) and Pi21388 (c) WT, AAAA-EER, RXLR-AAA and AAAA-AAA mRFP fusions were detected in the culture filtrate (CF) and mycelium (M) using αRFP. *P. infestans* histone H3 was detected using α H3 only in the M which indicates that the CF was not detectably contaminated by cellular material. Gold arrows indicate fusions cleaved at the EER motif, green arrows indicate fusions cleaved at the RXLR motif, and red arrows indicate fusions cleaved after the SP. Markers (kDa) are indicated on the left and protein loading is indicated by Ponceau stain (PS). Partial effector sequences containing the RXLR and EER motifs are shown below the PS. Immunoblots are representative of three independent biological replicates. (d) Distribution of the LC-MS/MS identified peptides within the sequence of Pi04314RXLR-AAA. Peptides were generated by trypsin in-gel digestion. Identified peptides are marked with red, predicted SP is shaded with grey, the RXLR motif is shaded with green, and the EER motif alanine replacement (AAA) is shaded with gold. (e) Graph shows the ratio of the number of times peptides immediately before (AEEENSIVAAAAAVETSEDEEER and GQLAGERAEEENSIVAAAAAVETSEDEEER, B) and after (DLLGLFAK and DLLGLFAKSK, A) the EER motif were detected. A two-tailed t-test with unequal variances indicates a statistically significant difference between the ratios of the lower and upper bands of Pi04314AAAA-EER (*: P < 0.05, 3 replicates).

To further confirm that the size difference between the effector RXLR and EER mutant mRFP fusions resulted from the different cleavage at the SP, RXLR or EER motifs, secreted Pi04314RXLR-AAA and Pi04314AAAA-EER mRFP fusions were immunoprecipitated from CF and analysed by LC-MS/MS. Peptides generated by trypsin in-gel digestion were only detected from the RXLR motif to the C-terminus of Pi04314RXLR-AAA (**Fig. 3d; Suppl Table S2e**), which suggests that the fusion was cleaved at the RXLR motif. As two obvious bands were observed in the CF of Pi04314AAAA-EER (**Fig. 3b; Suppl Fig. S7a 7b**), we excised and analysed them separately. For the upper band, peptides generated by trypsin in-gel digestion were detected immediately after the predicted SP to the C-terminus (**Suppl Fig. S7c; Suppl Table S2f**), which suggests that this band is the product from cleavage after the SP. Unexpectedly, similar sequence coverage as the upper band was observed with the lower band (**Suppl Fig. S7d; Suppl Table S2g**). Considering the close distance between the upper (≈42 kDa) and lower (≈36 kDa) bands and the inefficiency of SDS-PAGE to separate the two bands completely (**Suppl Fig. S7a**), it is likely that the detected peptides before the EER motif in the lower band were due to contamination of the upper band. Therefore, the excised upper band is a mixture of upper and lower bands dominated by the upper band while the excised lower band is the mixture dominated by the lower band. This can be reflected by the ratio of the number of times peptides immediately before (AEEENSIVAAAAAVETSEDEEER and GQLAGERAEEENSIVAAAAAVETSEDEEER) and after (DLLGLFAK and DLLGLFAKSK) the EER motif were detected. The upper band has a higher ratio compared with the lower band, and there is a statistically significant difference between them across replicates (**Fig. 3e; Suppl Table S2f, g**).

Since we were unable to separate the upper and lower bands of Pi04314AAAA-EER completely, we excised them together and digested with lys-C, which cuts only after lysine (K), instead of trypsin, which cuts after arginine (R) or K. Peptide coverage from immediately after the predicted SP to the C-terminus was observed (**Fig. 4a**), which is consistent with the trypsin digestion. A long peptide containing the alanine replacement of the RXLR motif, followed by the EER motif (GQLAGERAEEENSIVAAAAAVETSEDEEERDLLGLFAK), was detected multiple times (**Fig. 4b; Suppl Table S2h**), which further confirms that mutation of the RXLR motif to AAAA prevented cleavage at this site and the upper band resulted from the cleavage after the SP. Notably, multiple copies of a peptide from immediately after the EER motif (DLLGLFAK) were detected (**Fig. 4c; Suppl Table S2h**), which indicates that the lower band resulted from the endogenous cleavage after the R of the EER motif, since lys-C does not cut after R.

**Figure 4.**
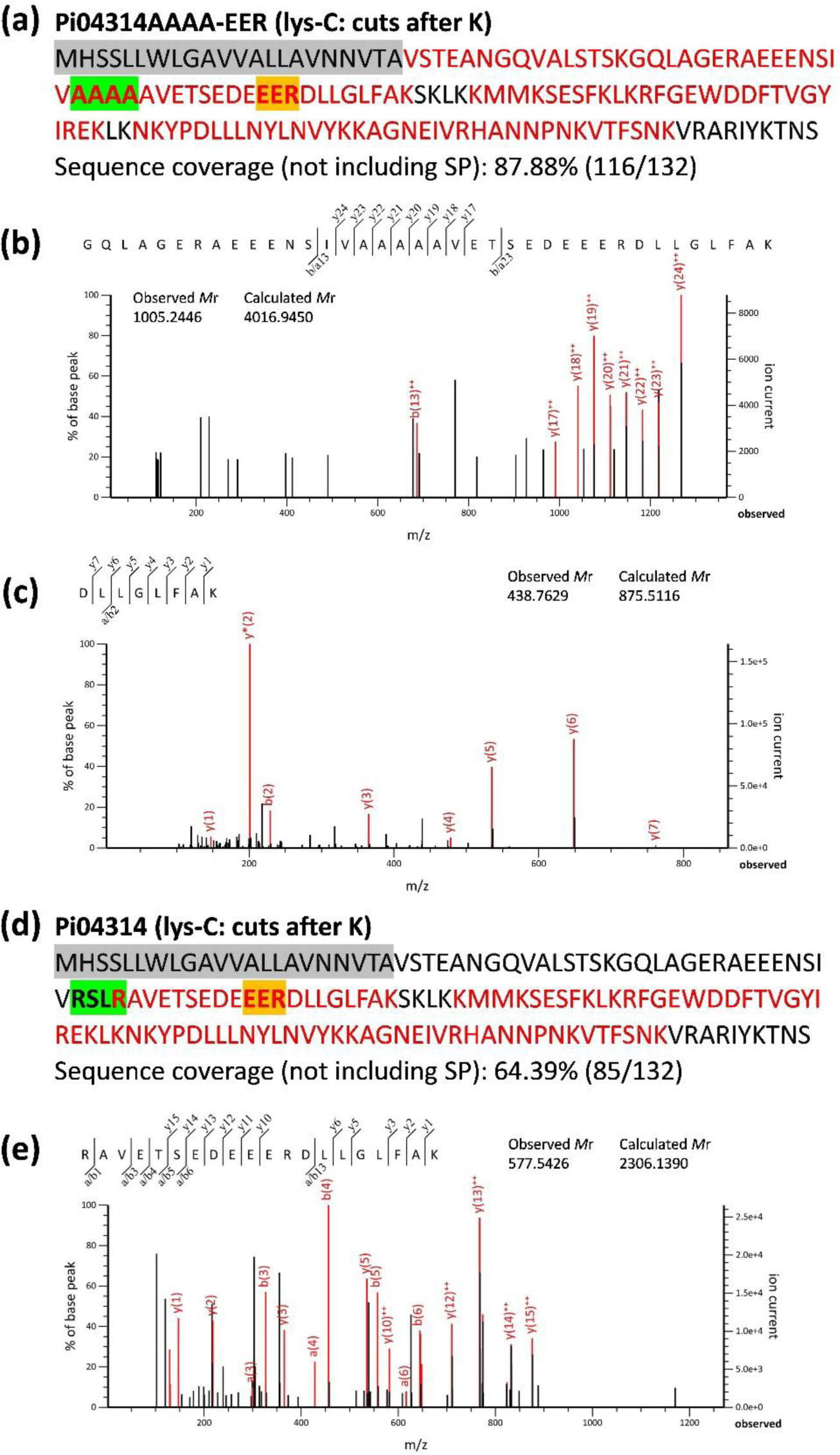
Cleavage of RXLR and EER motifs occurs after the leucine (L) and arginine (R) respectively. Distribution of the LC-MS/MS identified peptides within the sequences of Pi04314AAAA-EER (a) and WT (d). Peptides were generated by lys-C in-gel digestion. Identified peptides are marked with red, predicted SP is shaded with grey, the RXLR motif and its alanine replacement (AAAA) are shaded with green, and the EER motif is shaded with gold. (b) MS/MS fragmentation of the identified peptide GQLAGERAEEENSIVAAAAAVETSEDEEERDLLGLFAK containing the alanine replacement of the RXLR motif and the EER motif. (c) MS/MS fragmentation of the identified peptide DLLGLFAK starting immediately after the R of the EER motif. (e) MS/MS fragmentation of the identified peptide RAVETSEDEEERDLLGLFAK starting immediately after the L of the RXLR motif.

### The RXLR motif is cleaved after the leucine (L)

In the LC-MS/MS analysis of Pi04314-mRFP fusion protein peptides were detected from beyond the second R in the RXLR motif, which is consistent with the previous report about the cleavage of Avr3a (Wawra et al 2017). However, the detected peptides here and in the previous report were generated by in-gel digestion using trypsin (which cuts at the carboxyl side of R and K), which means the cleavage could be performed either by protease(s) of *P. infestans* before secretion or by trypsin during in-gel digestion. To further examine the precise site of cleavage at the RXLR motif, we first used the protease chymotrypsin (which cuts at the carboxyl side of F, L, W and Y) for in-gel digestion to test if a peptide starting immediately after the second R could still be detected. Peptides from Pi04314-mRFP digested with chymotrypsin represented sequences after the RXLR motif, which is consistent with trypsin digestion **(Suppl Fig. S8a)**. However, no peptides starting immediately after the second R were detected, whereas peptides starting immediately after the L (RAVETSEDEEERDLL) were detected **(Suppl Fig. S8b; Suppl Table S2i)**. This suggests that cleavage does not occur after the second R of the RXLR motif.

Since chymotrypsin cuts after L, we used the protease lys-C, which cuts at the carboxyl side of K, to account for the possibility that cleavage after the L occurred during in-gel digestion. Peptide coverage of Pi04314-mRFP generated by lys-C digestion was from the RXLR motif to the C-terminus **(Fig. 4d)**, which is consistent with the trypsin and chymotrypsin digestions. Similarly, multiple peptides starting immediately after the L (RAVETSEDEEERDLLGLFAK) were detected **(Fig. 4e; Suppl Table S2j)**, which confirms that cleavage in *P. infestans* at the RXLR motif occurs between the L and the second R. In addition, multiple copies of a peptide starting immediately after the EER motif (DLLGLFAK) were detected, which supports that endogenous cleavage also occurs after the R of the EER motif **(Suppl Table S2j)**.

Transformants expressing two further RFP-tagged RXLR-EER effectors were available in the laboratory; Pi04145 (Wang et al 2019) and Pi09160, which was detected in proteomics in endosomes isolated from infected plants (Wang et al 2023b). Western blotting of M and CF from cultures of these transformants showed bands consistent with cleavage at the RXLR and EER motifs (**Suppl Fig. S9a and S9b**). As with other such transformants, the effector fusions were secreted at haustoria during infection (**Suppl FigS9c and S9d**). These observations support the hypothesis that cleavage at RXLR and EER motifs is a general rule.

### Efficient cleavage of RXLR and EER motifs does not occur during secretion from plant cells

To test whether RXLR-only and RXLR-EER effectors of *P. infestans* would be similarly proteolytically processed during secretion from other eukaryotic cells we expressed full-length (including SP) versions of Pi09216-mRFP (RXLR-only), Pi04314-mRFP and Pi21388-mRFP (RXLR-EER) in *N. benthamiana* cells. As controls, we also expressed non-secreted truncated versions from the end of the RXLR to the C-terminus (Pi09216_38-175_-mRFP, Pi04314_57-154_-mRFP and Pi21388_55-152_-mRFP) and from the end of the EER to the C-terminus (Pi04314_68-154_-mRFP and Pi21388_73-152_-mRFP) in *N. benthamiana*. The truncated versions were each of an expected size and revealed subcellular localisations observed previously (**Fig. 5)**; Pi09216-mRFP at small puncta shown previously to include mitochondria (Wang et al 2019) and Pi04314-mRFP and Pi21388-mRFP localised primarily in host nuclei and nucleoli (Boevink et al 2016; Wang S et al 2019; Wang H et al 2023b). By contrast, the full-length effector fusions with SP were each localised at the cell periphery, consistent with them being secreted by the host cells (**Fig. 5a, 5c, 5e**). Interestingly, the full length Pi09216-mRFP was primarily of a size predicted from processing at the SP and was only very weakly cleaved to generate a fusion protein of the size expected from RXLR cleavage (**Fig. 5b**). The full-length versions of Pi04314-mRFP and Pi21388-mRFP again each showed fusion proteins primarily processed after the SP, faint bands of a size consistent with weak cleavage at the RXLR motif, and no evidence of processing at the EER motif (**Fig. 5d, 5f)**.

**Figure 5.**
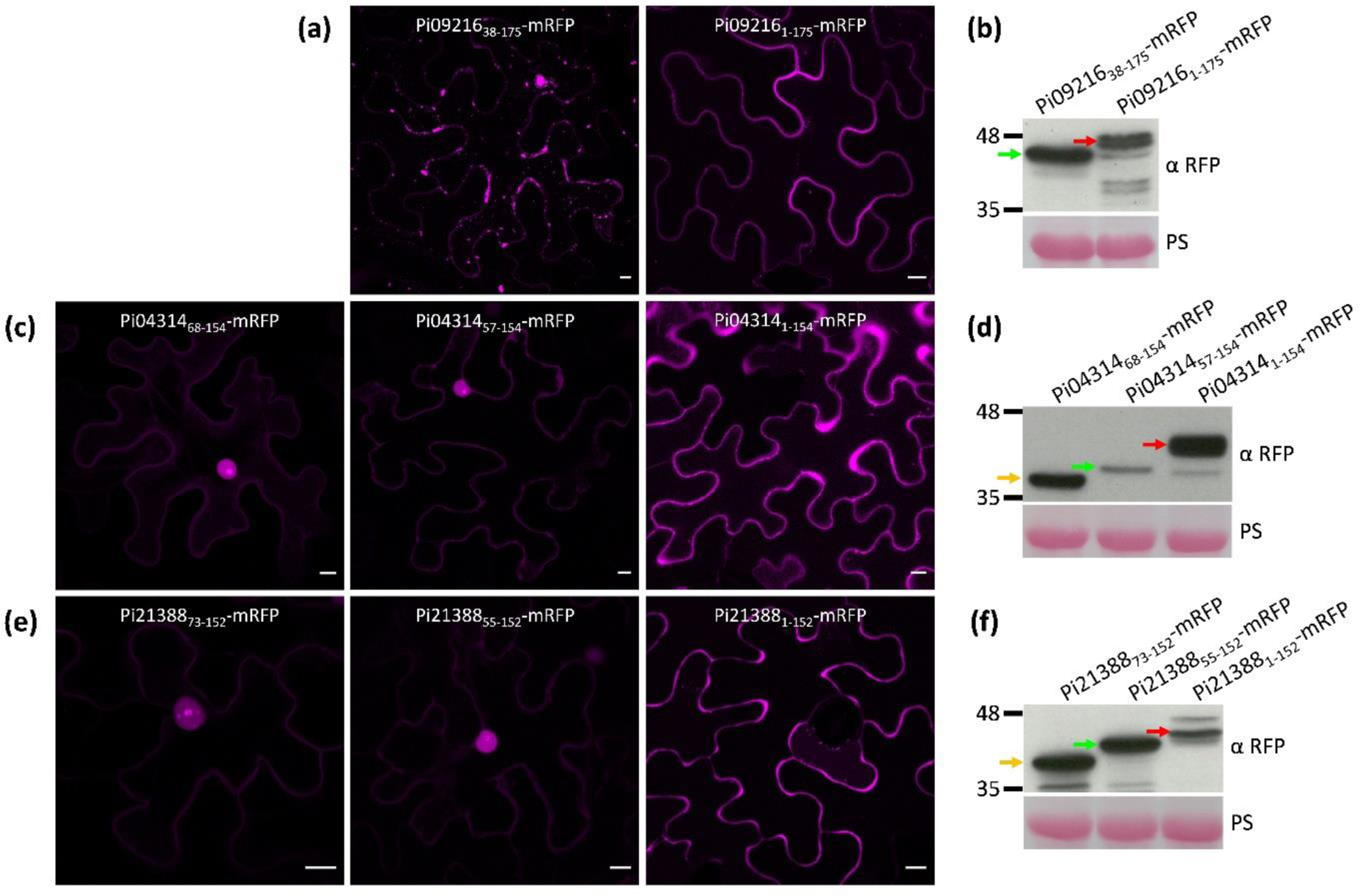
Efficient cleavage of RXLR and EER motifs does not occur during secretion from the plant. Confocal images of *Agrobacterium*-mediated transient overexpression of Pi09216_38-175_-mRFP (a, left), Pi09216_1-175_-mRFP (a, right), Pi04314_68-154_-mRFP (c, left), Pi04314_57-154_-mRFP (c, middle), Pi04314_1-154_-mRFP (c, right), Pi21388_73-152_-mRFP (e, left), Pi21388_55-152_-mRFP (e, middle) and Pi21388_1-152_-mRFP (e, right) in *Nicotiana benthamiana* cells. Images were taken 2 days post infiltration. Scale bars represent 10 μm. Expression of Pi09216_38-175_-mRFP and Pi09216_1-175_-mRFP (b), Pi04314_68-154_-mRFP, Pi04314_57-154_-mRFP and Pi04314_1-154_-mRFP (d), and Pi21388_73-152_-mRFP, Pi21388_55-152_-mRFP and Pi21388_1-152_-mRFP (f) fusions was confirmed by immunoblotting using α RFP. Markers (kDa) are indicated on the left and protein loading is indicated by Ponceau stain (PS). Red, green and gold arrows indicate effector-mRFP fusions starting after the SP, RXLR motif, and EER motif, respectively.

### Cleavage of the RXLR motif is related to its position

There are two RXLR motifs within the sequence of effector Pi21388; one is in the predicted unstructured region of the N-terminus and the other is located within a short alpha helix in the C-terminal domain **(Fig. 6a)**. According to immunoblots detecting WT Pi21388-mRFP in M and CF samples **(Fig. 2c**, **Fig. 3c)**, only the N-terminal RXLR motif was cleaved, which suggests that cleavage of RXLR motif could be related to its location within the sequence. To test this hypothesis, an extra RXLR motif was introduced into the sequence of WT Pi04314 and RXLR mutant Pi04314AAAA-EER by mutating the original amino acid residues KSKL, which are 7 residues downstream of the EER motif and located within a predicted alpha helix, to RSLR **(Fig. 6b, 6c)**. Transformants overexpressing Pi04314RXLR-EER-RXLR-mRFP or Pi04314AAAA-EER-RXLR-mRFP were generated and the secretion of these mRFP fusions at haustoria during infection was confirmed by confocal microscopy (**Suppl Fig. S10a)**. The CF and M of these transformants grown *in vitro* were used for protein precipitation and extraction, respectively, and samples were analysed by immunoblotting along with samples from the Pi04314 WT and AAAA-EER transformants. The number and size of Pi04314RXLR-EER-RXLR-mRFP protein bands are the same as WT Pi04314-mRFP **(Fig. 6d)**, which suggests that the introduced RXLR motif is not cleaved. Moreover, the number and size of Pi04314AAAA-EER-RXLR-mRFP protein bands is also the same as Pi04314 AAAA-EER-mRFP **(Fig. 6d)**, which indicates that the introduced RXLR motif is not proteolytically cleaved. Similar results were observed in an additional two replicates (**Suppl Fig. S10b)**. Together, we concluded that cleavage of the RXLR motif is related to its position.

**Figure 6.**
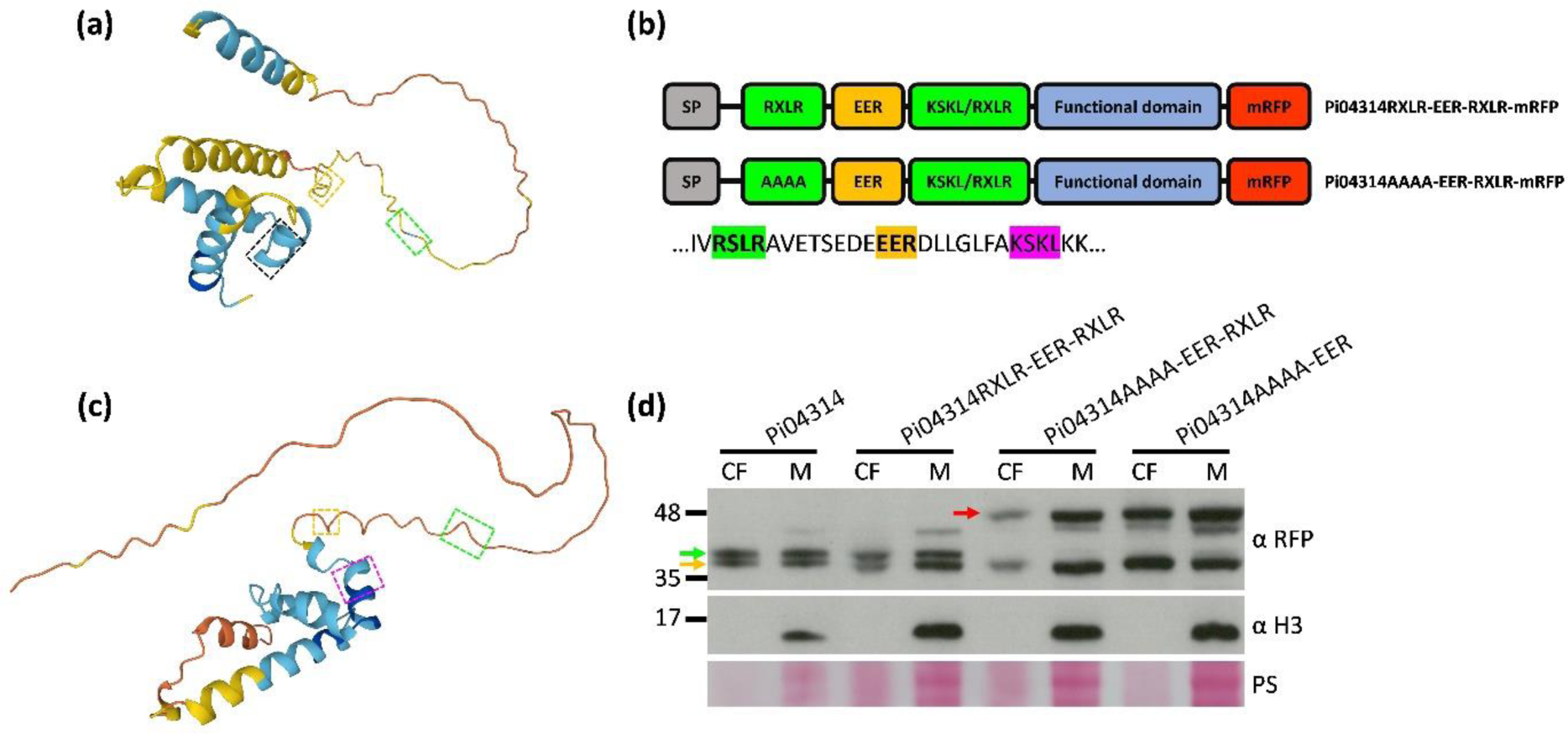
Cleavage of the RXLR motif is related to its position. Predicted structures of Pi21388 (a) and Pi04314 (c) obtained from the AlphaFold Protein Structure Database (https://alphafold.ebi.ac.uk/). RXLR motifs in the N-terminus are enclosed by green dashed rectangles, EER motifs are enclosed by gold dashed rectangles, another RXLR motif of Pi21388 in the C-terminus is enclosed by the black dashed rectangle, and the KSKL amino acids of Pi04314 are enclosed by the purple dashed rectangle. (b) Schematic representation of Pi04314RXLR-EER-RXLR and Pi04314AAAA-EER-RXLR mRFP fusions. Partial Pi04314 sequence containing the RXLR (shaded with green), EER (shaded with gold) motifs and the KSKL amino acids (shaded with purple) is shown below. (d) Pi04314 WT, RXLR-EER-RXLR, AAAA-EER-RXLR and AAA-EER mRFP fusions were detected in the culture filtrate (CF) and mycelium (M) using α RFP. *P. infestans* histone H3 was detected using α H3 only in the M which indicates that the CF was not detectably contaminated by cellular material. Gold arrow indicates fusions cleaved at the EER motif, green arrow indicates fusions cleaved at the RXLR motif, and red arrow indicates fusions cleaved after the SP. Markers (kDa) are indicated on the left and protein loading is indicated by Ponceau stain (PS). Immunoblot is representative of three independent biological replicates.

## Discussion

In this study it was found that the RXLR and EER motifs are two independent proteolytic cleavage sites within *P. infestans* RXLR effectors that are processed prior to secretion. These processing steps appears to be shared generally by different RXLR effectors. Based on the immunoblotting and proteomics analyses, a model of how RXLR effectors are processed before secretion was proposed **(Fig. 7)**.

**Figure 7.**
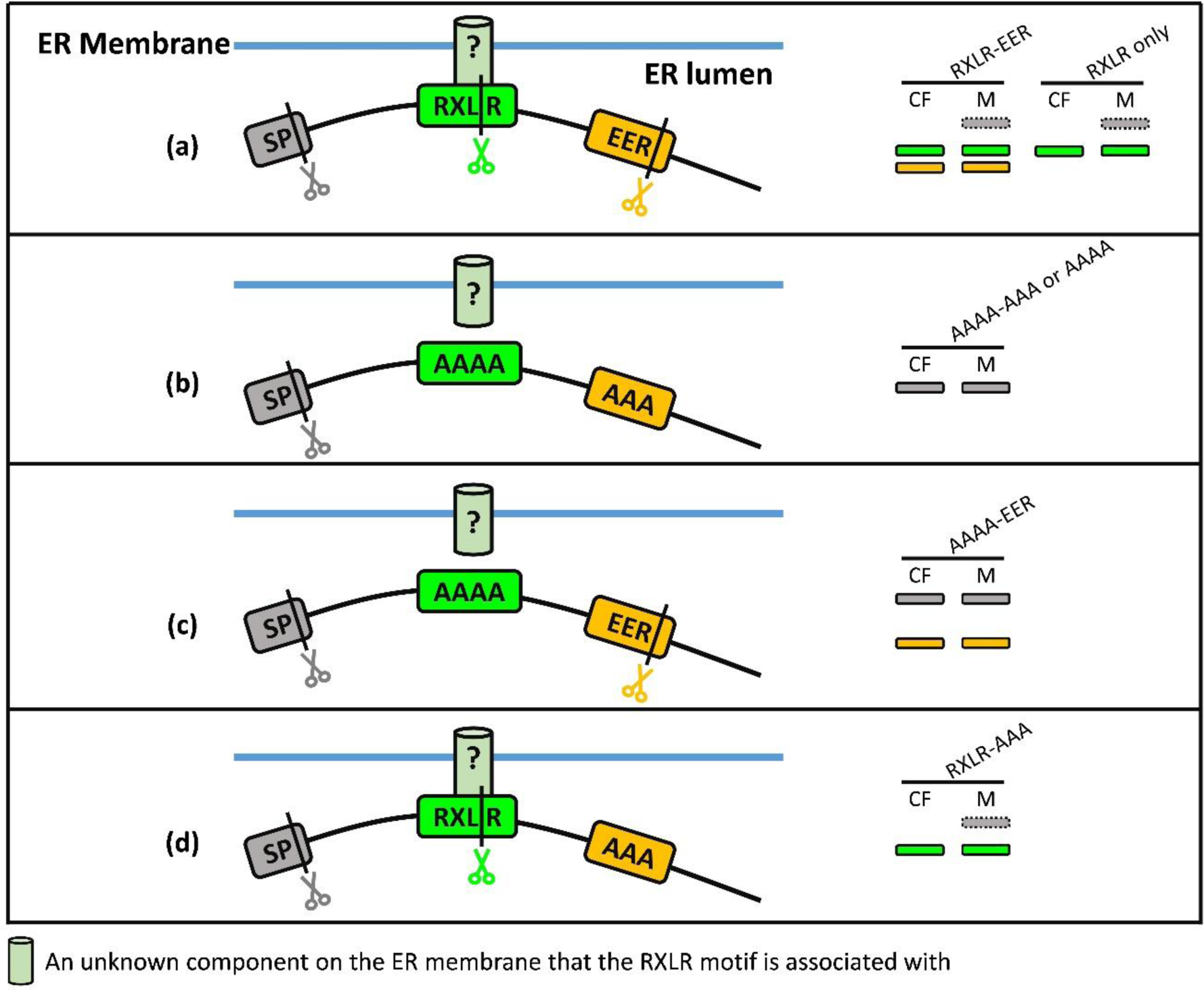
A model for RXLR(-EER) effector cleavage. SP directs effector into the endoplasmic reticulum (ER), where SP is cleaved. The RXLR motif is associated with an unknown component (indicated as light green cylinders), potentially at the ER membrane (indicated as blue straight lines). (a) WT RXLR-EER effectors are cleaved at the RXLR and EER motifs, resulting in the presence of two lower bands in the CF and M (cleaved at the RXLR and EER motifs, represented by green and gold rectangles), or one lower band for RXLR-only effectors. Some effectors are incompletely cleaved resulting in the presence of a higher band (cleaved after the SP, represented by the grey dashed rectangles) which is retained in the M due to the association with an unknown component. (b) When both the RXLR and EER motifs are mutated, the association between the RXLR and the unknown component and the cleavage events are abolished, resulting in only one higher band in both the CF and M (cleaved after the SP, represented grey rectangles). (c) When only mutating the RXLR motif, the association between the RXLR and the unknown component is abolished, the cleavage at the RXLR is disrupted but the cleavage at the EER still occurs, resulting in the presence of one higher band (cleaved after the SP, grey rectangles) and one lower band (cleaved at the EER, gold rectangles). (d) When only mutating the EER motif, the RXLR is still associated with the unknown component, the cleavage at the RXLR still occurs, but the cleavage at the EER is disrupted, resulting in the presence of one lower band (cleaved at the RXLR, green rectangles). Incompletely cleaved effectors (cleaved after the SP, grey dashed rectangle) are retained in the M. Black vertical lines indicate the cleavage sites of the SP, RXLR and EER cleavage. The grey scissors represent the signal peptidase that cleaves SP, the green scissors represent the protease that cleaves at the RXLR, and the gold scissors represent the protease that cleaves at the EER.

Proteolytic cleavage at the RXLR motif was observed in the effector Avr3a and was reported to occur after the second R (Wawra et al. 2017). In this study, we also observed cleavage at the RXLR motif in several RXLR effectors, thus demonstrating that this is likely a general processing step. However, by using different protein digestion enzymes for mass spectrometry analysis we found that the cleavage occurs after the L, not the second R for effector Pi04314. Whilst this needs to be confirmed for other RXLR effectors, cleavage after the L is consistent with processing of the RXLXE/D/Q motif of *P. falciparum* PEXEL effectors (Boddey et al., 2010; Russo et al., 2010; Marapana et al 2018). Variations in sequence of the RXLR motif have been observed in effectors of downy mildews, including GXLR/RXLT in *Hyaloperonospora arabidopsidis* (Baxter et al., 2010), RXLK/Q/D in *Bremia lactucae* (Stassen et al., 2012), RXLK/Q/E in *Plasmopara halstedii* (Sharma et al., 2015), and RXLK/Q in white rusts such as *Albugo candida* and *A. laibachii* (Kemen et al., 2011; Links et al., 2011). These variations occur at the Rs of the RXLR motif but not the L, which supports the importance of this residue as a conserved component of the cleavage site. This fits with the observation that cleavage occurs immediately after this residue, rather than the variable second R in the motif. Further studies are needed to confirm the conservation of this processing site across the oomycetes.

The position of the RXLR motif, relative to the SP, has been postulated to be restricted (Win et al 2007). Moreover, we observed that the cleaved RXLR motifs in Pi21388/AvrBlb1 and Pi04314 were in predicted disordered regions. Indeed, we observed that a second naturally occurring RXLR motif in Avrblb1, which was not cleaved, was located closely following the EER motif in a structured region of the effector (**Fig. 6**). This prompted us to introduce a second RXLR motif into a similar location in Pi04314, again in a predicted structured region. This RXLR motif was also not cleaved (**Fig. 6**), supporting a hypothesis that the cleaved RXLR motifs are positioned in disordered regions within 30-40 amino acids of the SP to facilitate efficient proteolytic processing.

Cleavage at the EER motif was demonstrated for the first time in this study, but the biological significance of this processing event is unknown. Studies have shown that mutation at the EER motif disables translocation of Avr3a (Whisson et al. 2007), so it may be possible that EER cleavage is involved in the trafficking of RXLR effectors. Indeed, some effectors from *B. lactucae* and *H. arabidopsidis* which are predicted to possess the conserved WY structural fold in the effector domain, appear to only possess the EER motif. They are nevertheless postulated to be translocated and recognized by host resistance (R) proteins (Bailey et al., 2011; Stassen et al., 2012; Wood et al., 2020). However, some RXLR effectors that only have the RXLR and no EER motif can also be translocated and recognized, such as those from the Avrblb2 family (Haas, et al., 2009; Oh et al., 2009). This is consistent with a hypothesis that there may be a functional overlap between the RXLR and EER motifs in mediating RXLR effector translocation.

Identifying proteases that are involved in critical cellular processes is of great importance, as they can be potential targets for anti-oomycete drug development. Wawra et al. (2017) tried to identify proteases that are responsible for the cleavage of Avr3a. Eleven pepsin-like aspartic proteinases (APs) of *P. infestans* were tested as some of them are homologous to *P. falciparum* plasmepsin V (PMV) which cleaves PEXEL effectors. However, no cleavage activity was detected with these candidates (Wawra et al., 2017). Schoina et al. (2019) reported that PiAP10 and PiAP12 have weak cleavage activity towards *P. infestans* RXLR effector AVR4 *in vitro*. Further studies are needed to identify and test protease candidates. Protein cleavage usually involves transient interaction between the protease and its substrate. TurboID-based proximity labelling (PL) has been demonstrated to be useful for the identification of weakly and transiently interacting proteins (Branon et al., 2019; Zhang et al., 2019). It could be adopted to identify potential protease candidates that are responsible for cleavage of the RXLR or EER motifs. Cleavage of RXLR effectors likely occurs in the ER, so we may be able to narrow down candidates based on (i) possession of a SP for ER entry, and (ii) possession of an ER-retention signal or a transmembrane domain (TMD) for ER retention. We demonstrated that plant proteases can only poorly cleave the RXLR motif and are incapable of cleaving the EER motif when full-length effectors are secreted from *N. benthamiana* cells **(Fig. 5b, d, f)**. This raises the potential that co-expression of RXLR effectors and candidate *Phytophthora* proteases in the plant ER may provide an initial screen of protease candidates.

For RXLR-only effector Pi04097 and RXLE-EER effector Pi22926, there was a higher molecular weight band exclusively in mycelium samples but not the culture filtrate (CF) of a size consistent with cleavage only at the signal peptide (SP) **(Figs 1c, S5c)**. This suggests that some WT effectors, following cleavage at the SP but prior to RXLR cleavage, were retained in the pathogen cell. One hypothesis is that SP cleavage occurs prior to RXLR processing and the RXLR motif may be associated with an unknown component that anchors the effectors to the ER luminal membrane (**Fig. 7)**. Cleavage at the RXLR motif subsequently releases effectors for secretion. Sometimes, this higher mycelium-exclusive band can also be detected in the mycelium of Pi04314 WT **(Figs 6d, S5a, S10b)**, but it is much fainter compared with Pi22926 and Pi04097 WT. We speculate that the intensity of this band is related to the efficiency of proteases releasing effectors from the association with the unknown retention component.

One possibility for the unknown component that the RXLR motif could associate with is phosphatidylinositol monophosphates (PIPs). Kale et al. (2010) reported that the RXLR motif can specifically bind to PI3P on the outer surface of plant (soybean root cell) and human (A549 cell) cell plasma membrane, and the binding mediates the entry of RXLR effectors into host and non-host cells. Bhattacharjee et al. (2012) found that the binding between the RXLR motif of the *P. infestans* RXLR effector Nuk10 (PITG_15287) and PI3P on the ER luminal membrane of *P. falciparum* mediates export of the effector. However, RXLR-PI3P binding is highly controversial (Ellis & Dodds, 2011; Yaeno et al., 2011; Wawra et al., 2012; Boddey et al., 2016). An alternative candidate for the unknown binding component could be the ER translocon complex. Marapana et al. (2018) demonstrated that the recessed SPs of *P. falciparum* PEXEL effectors could act as uncleaved signal anchors to retain the full-length proteins on the ER membrane before the PEXEL cleavage by interacting with the special ER translocon complex that involves the protease PMV. Considering the phylogenetic relatedness between *P. infestans* and *P. falciparum*, it is possible that the RXLR motif may have a similar role as the recessed SPs of PEXEL effectors to anchor effectors on the ER membrane by interacting with a specialised ER translocon complex.

In conclusion, we reveal that both RXLR and EER motifs are sites for proteolytic cleavage. This is of significant interest as effectors possessing only the RXLR or EER motif have been observed, raising the possibility that they have a redundant or overlapping role in effector trafficking. This work thus establishes the need to determine the precise biological functions of each motif and to identify the proteases that cleave them. Such knowledge would provide biochemical or genetic targets for control of multiple diseases that threaten food, energy and environmental security.

## Acknowledgements

We thank colleagues in the Division of Plant Sciences, University of Dundee, and the Cell and Molecular Sciences Department, James Hutton Institute, for helpful discussions and insightful suggestions throughout the course of this research. This work was financially supported by the Biotechnology and Biological Sciences Research Council grant BB/S003096/1; European Research Council (ERC)-Advanced grant PathEVome (787764); and Scottish Government Rural and Environment Science and Analytical Services Division (RESAS) JHI B1-1. WW and LX are supported by the China Scholarship Council (CSC).

## Author Contributions

LX, SCW, PCB and PRJB designed the work; LX, SW, HW, WW, and LW, performed the research; LX, SCW, PCB and PRJB analysed and interpreted data; LX prepared the Figures; SCW, PCB and PRJB secured the funding and supervised the work; LX and PRJB wrote the manuscript with input from all co-authors.

