## Supplementary Figures for "Proteolytic processing of both RXLR and EER motifs in oomycete effectors"

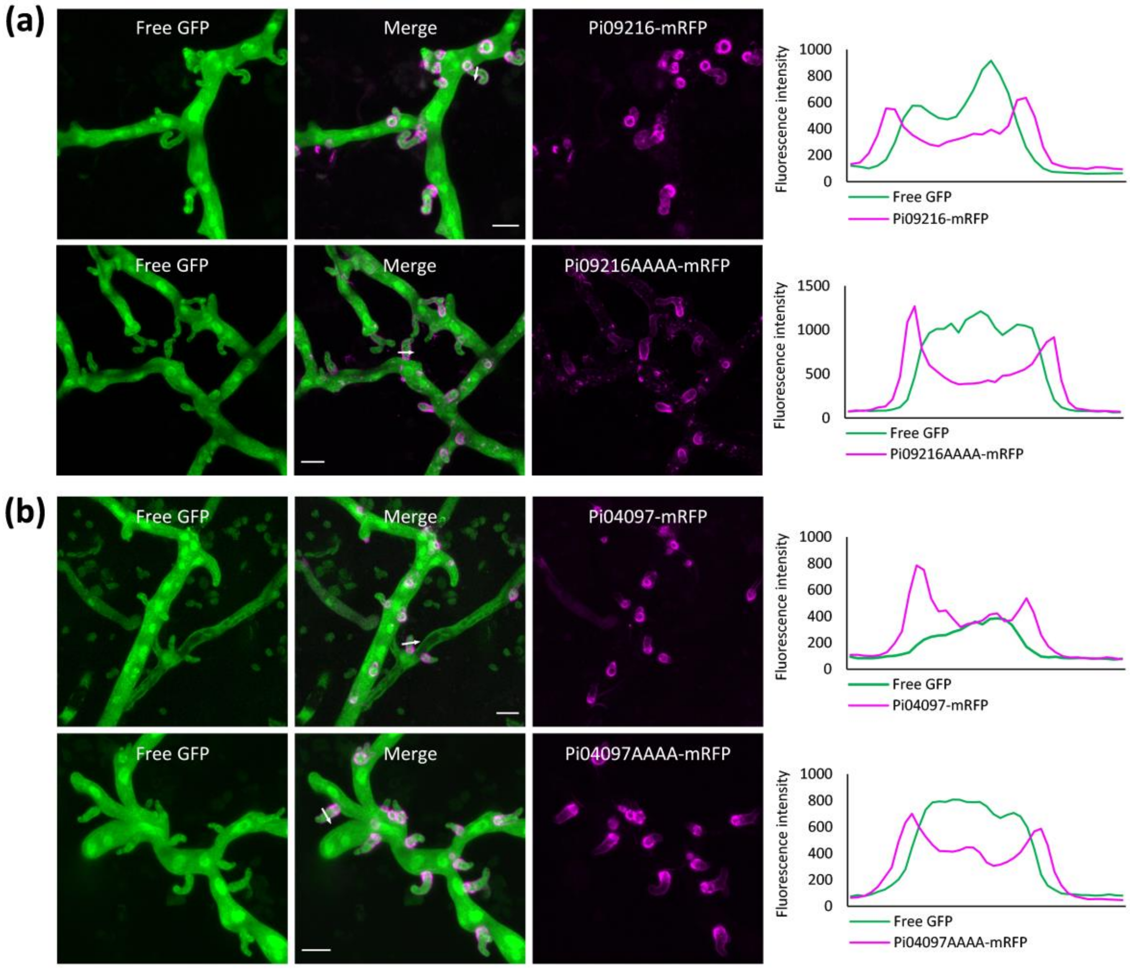


**Figure S1 Secretion of RXLR-only effector wild type (WT) and RXLR mutant (AAAA) *in planta*.** Confocal images of *P. infestans* hyphae and haustoria of Pi09216 WT (a, upper panel), Pi09216AAAA (a, lower panel), Pi04097 WT (b, upper panel) and Pi04097AAAA (b, lower panel) transformants infecting *N. benthamiana* leaves. Effector WT and AAAA mRFP fusions were secreted from haustoria and accumulated at the periphery of haustoria (indicated by magenta). Hyphae and haustoria were visualized by the co-expressed cytoplasmic GFP. Images are projections of confocal z-series. Scale bars represent 10 µm. White arrows indicate the lines used for the corresponding fluorescence intensity profiles shown on the right of the images.

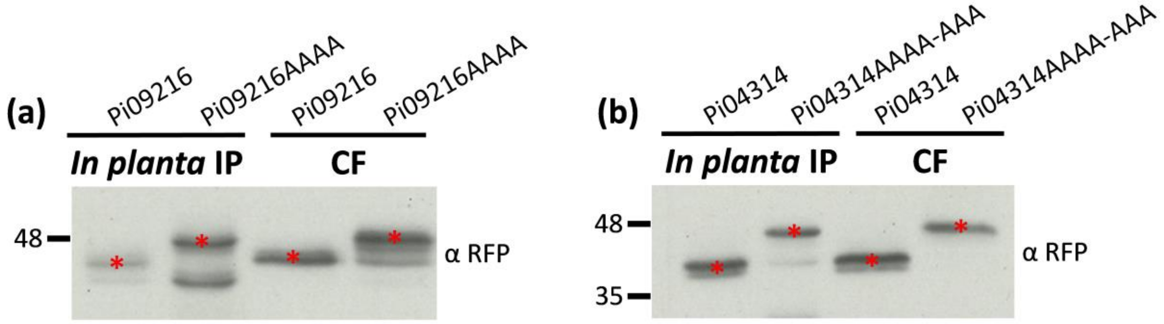


**Figure S2 Cleavage of Pi09216 and Pi04314 WT during infection.** Cleaved Pi09216 (a) and Pi04314 (b) WT mRFP fusions and uncleaved Pi09216AAAA (a) and Pi04314 RXLR-EER mutant (Pi04314AAAA-AAA) (b) mRFP fusions were detected in the immunoprecipitation (IP) samples from infected *N. benthamiana* leaves and the culture filtrate (CF) using αRFP. Bands with expected sizes are indicated by red asterisk and markers (kDa) are indicated on the left.

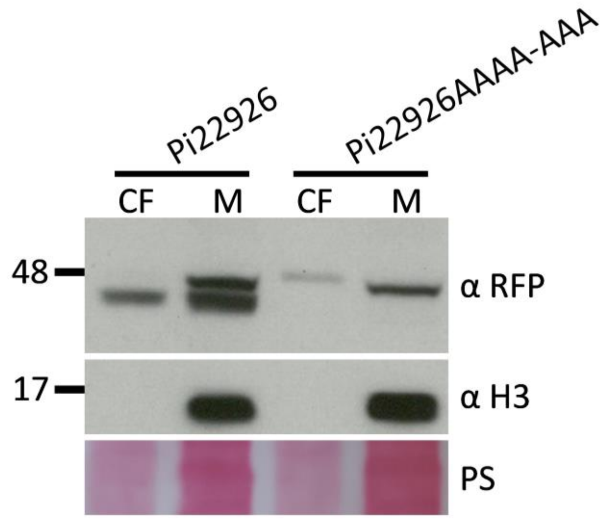


**Figure S3 Cleavage of RXLR-EER effector Pi22926.** Pi22926 WT and AAAA-AAA mRFP fusions were detected in the culture filtrate (CF) and mycelium (M) using αRFP. *P. infestans* histone H3 was detected using α H3 only in the M which indicates that the CF was not detectably contaminated by cellular material. Markers (kDa) are indicated on the left and protein loading is indicated by Ponceau stain (PS).

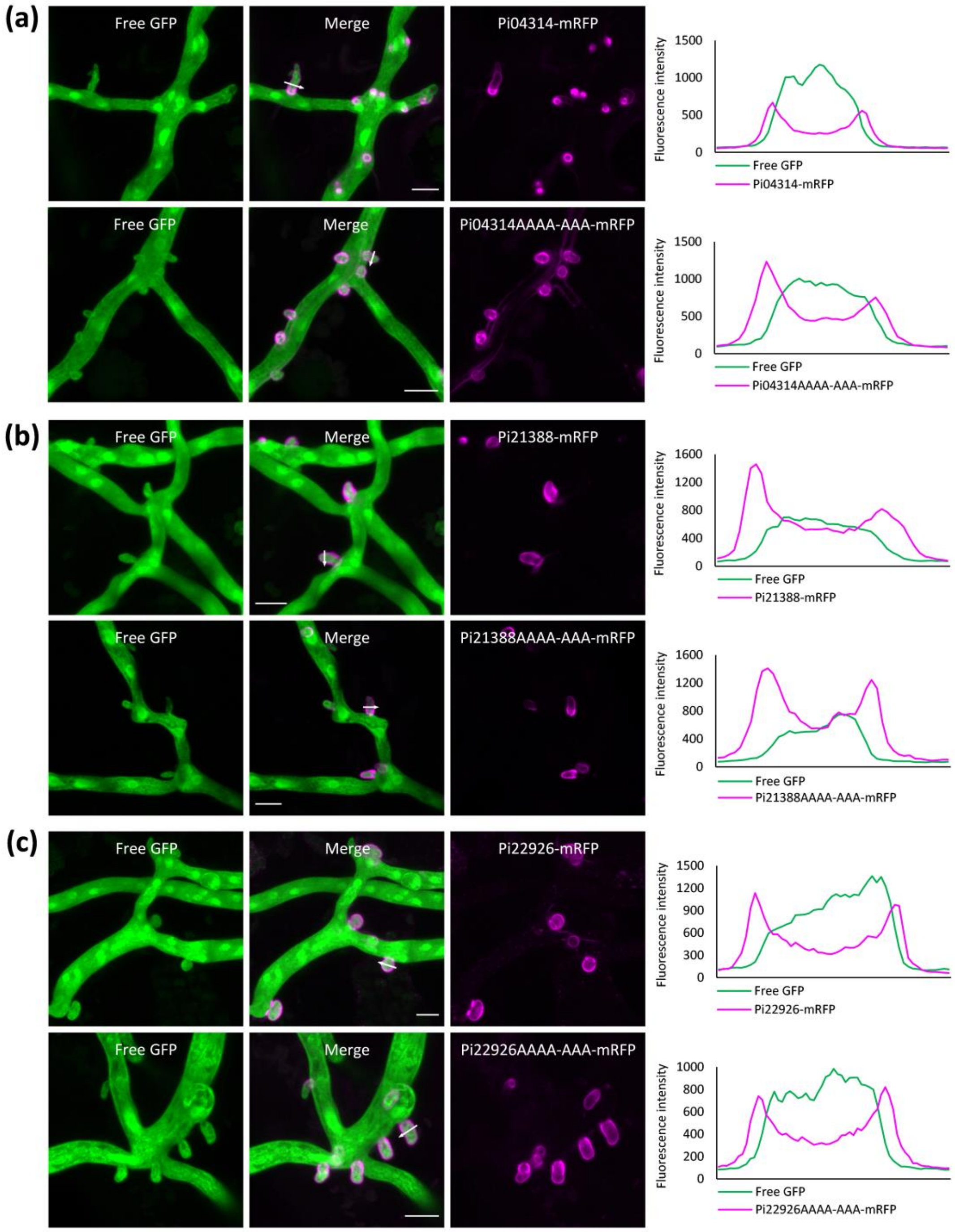


**Figure S4. Secretion of RXLR-EER effector WT and AAAA-AAA *in planta*.** Confocal images of *P. infestans* hyphae and haustoria of Pi04314 WT (a, upper panel), Pi04314AAAA-AAA (a, lower panel), Pi21388 WT (b, upper panel), Pi21388AAAA-AAA (b, lower panel), Pi22926 WT (c, upper panel) and Pi22926AAAA-AAA (c, lower panel) transformants infecting *N. benthamiana* leaves. Effector WT and AAAA-AAA mRFP fusions were secreted from haustoria and accumulated at the periphery of haustoria (indicated by magenta). Hyphae and haustoria were visualized by the co-expressed cytoplasmic GFP. Images are projections of confocal z-series. Scale bars represent 10 µm. White arrows indicate the lines used for the corresponding fluorescence intensity profiles shown on the right of the images.

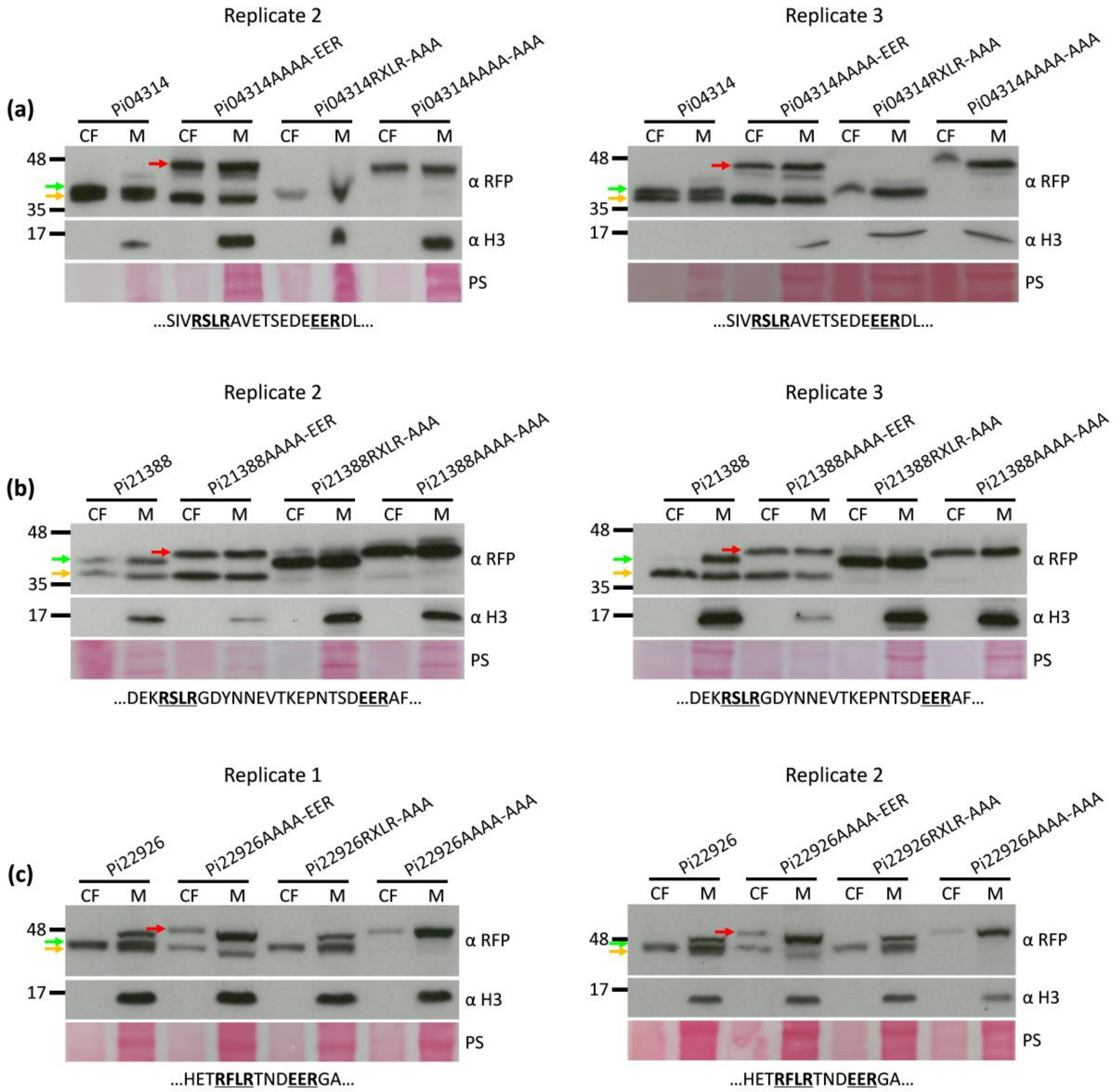


**Figure S5. Detection of RXLR-EER effector WT and mutants by immunoblotting.** Pi04314 (a), Pi21388 (b) and Pi22926 (c) WT, AAAA-EER, RXLR-AAA and AAAA-AAA mRFP fusions were detected in the culture filtrate (CF) and mycelium (M) using αRFP in two independent replicates. *P. infestans* histone H3 was detected using α H3 only in the M which indicates that the CF was not detectably contaminated by cellular material. Gold arrows indicate fusions cleaved at the EER motif, green arrows indicate fusions cleaved at the RXLR motif, and red arrows indicate fusions cleaved after the SP. Markers (kDa) are indicated on the left and protein loading is indicated by Ponceau stain (PS). Partial effector sequences containing the RXLR and EER motifs are shown below the PS.

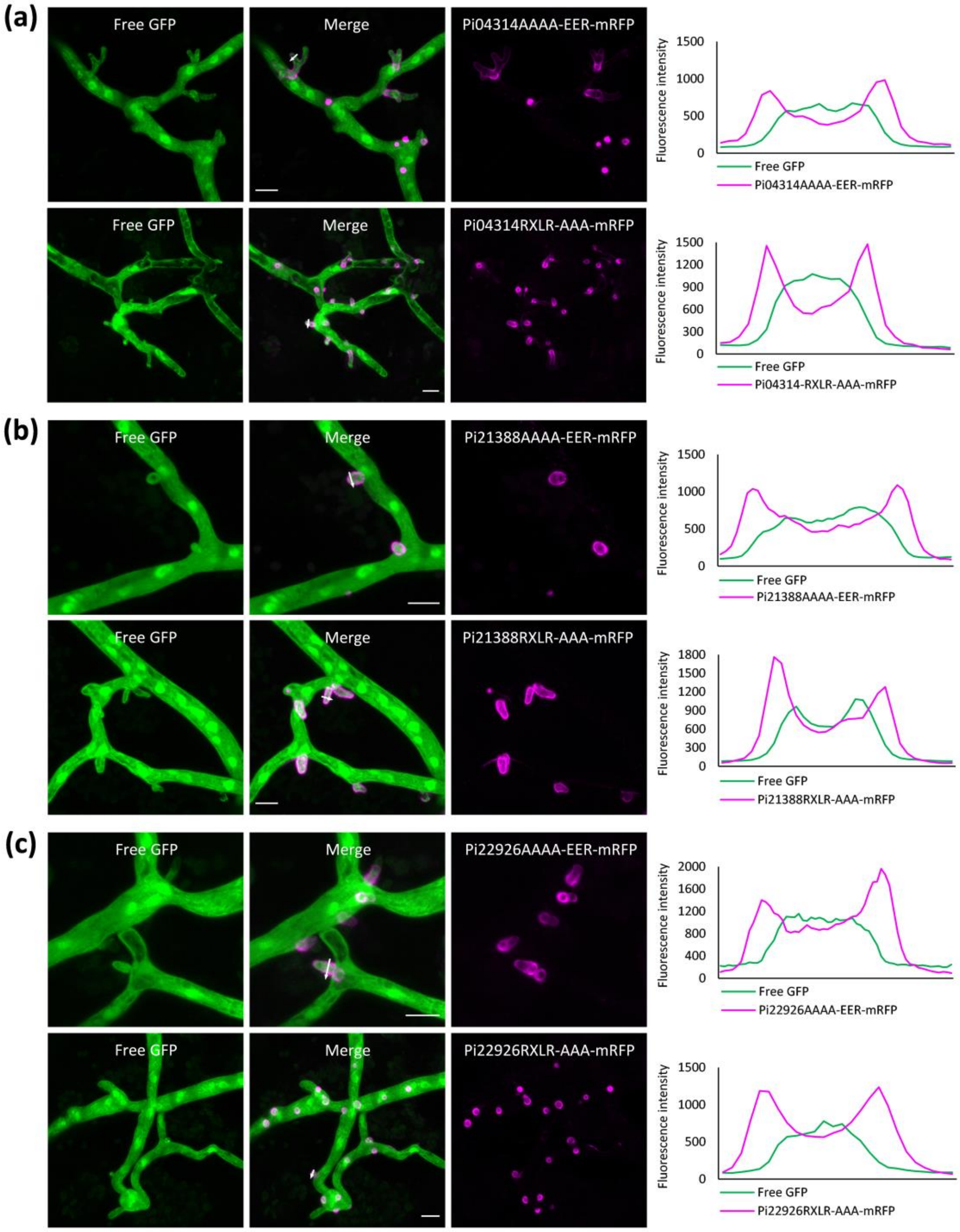


**Figure S6 Secretion of RXLR-EER effector AAAA-EER and RXLR-AAA *in planta*.** Confocal images of *P. infestans* hyphae and haustoria of Pi04314AAAA-EER (a, upper panel), Pi04314RXLR-AAA (a, lower panel), Pi21388AAAA-EER (b, upper panel), Pi21388RXLR-AAA (b, lower panel), Pi22926AAAA-EER (c, upper panel) and Pi22926RXLR-AAA (c, lower panel) transformants infecting *N. benthamiana* leaves. Effector AAAA-EER and RXLR-AAA mRFP fusions were secreted from haustoria and accumulated at the periphery of haustoria (indicated by magenta). Hyphae and haustoria were visualized by the co-expressed cytoplasmic GFP. Images are projections of confocal z-series. Scale bars represent 10 µm. White arrows indicate the lines used for the corresponding fluorescence intensity profiles shown on the right of the images.

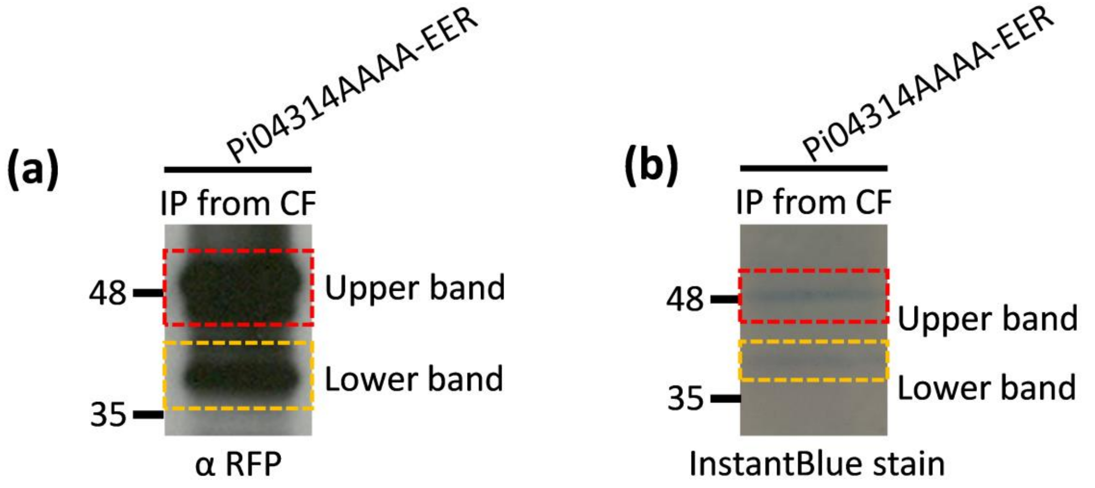


(c) **Pi04314AAAA-EER upper band (trypsin: cuts after K/R)**

MHSSLLWLGAVVALLAVNNVTAVSTEANGQVALSTSKGQLAGERAEEENSI
VAAAAAVETSEDEEERDLLGLFAKSKLKKMMKSESFKLKRFGEWDDFTVGY
IREKLKNKYPDLLLNYLNVYKKAGNEIVRHANNPNKVTFSNKVRARIYKTNS
Sequence coverage (not including SP): 90.15% (119/132)

(d) **Pi04314AAAA-EER lower band (trypsin: cuts after K/R)**

MHSSLLWLGAVVALLAVNNVTAVSTEANGQVALSTSKGQLAGERAEEENSI
VAAAAAVETSEDEEERDLLGLFAKSKLKKMMKSESFKLKRFGEWDDFTVGY
IREKLKNKYPDLLLNYLNVYKKAGNEIVRHANNPNKVTFSNKVRARIYKTNS
Sequence coverage (not including SP): 90.15% (119/132)

**Figure S7. LC-MS/MS analysis of Pi04314AAAA-EER upper and lower bands.** Immunoprecipitated Pi04314AAAA-EER mRFP fusion from the CF was detected using αRFP (a) and InstantBlue stain (b). Upper bands (cleaved after the SP) are enclosed by red dashed rectangles, and lower bands (cleaved at the EER motif) are enclosed by gold dashed rectangles. Markers (kDa) are indicated on the left. Distribution of the LC-MS/MS identified peptides from the upper (c) and lower (d) bands within the sequence of Pi04314AAAA-EER. Peptides were generated by trypsin in-gel digestion. Sequence coverage by identified peptides is marked with red, predicted SP is shaded with grey, the RXLR motif alanine replacement (AAAA) is shaded with green, and the EER motif is shaded with gold. K: lysine, R: arginine.

**(a) Pi04314 (chymotrypsin: cuts after F/L/W/Y)**

MHSSLLWLGAVVALLAVNNVTAVSTEANGQVALSTSKGQLAGERAEEENSI
VRSLRAVETSEDEEERDLLGLFAKSKLKKMMKSESFKLKRFGEWDDFTVGYI
REKLKNKYPDLLLNYLNVYKKAGNEIVRHANNPNKVTFSNKVRARIYKTNS
Sequence coverage (not including SP): 60.61% (80/132)

**(b)**

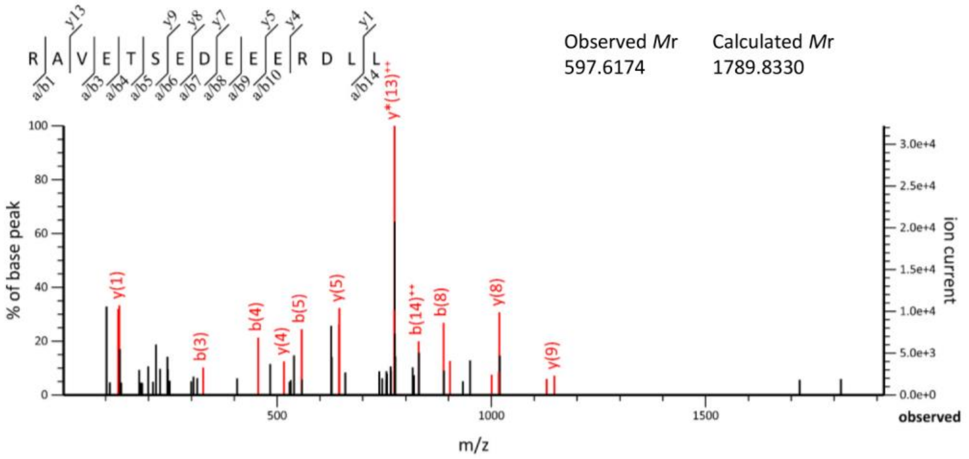


**Figure S8. Additional evidence for the cleavage after L in the RXLR motif.** (a) Distribution of the LC-MS/MS identified peptides within the sequences of Pi04314 WT. Peptides were generated by chymotrypsin in-gel digestion. Sequence coverage by identified peptides is marked with red, predicted SP is shaded with grey, the RXLR motif is shaded with green, and the EER motif is shaded with gold. F: phenylalanine, L: leucine, W: tryptophan, Y: tyrosine. (b) MS/MS fragmentation of the identified peptide RAVETSEDEEERDLL starting immediately after the L of the RXLR motif.

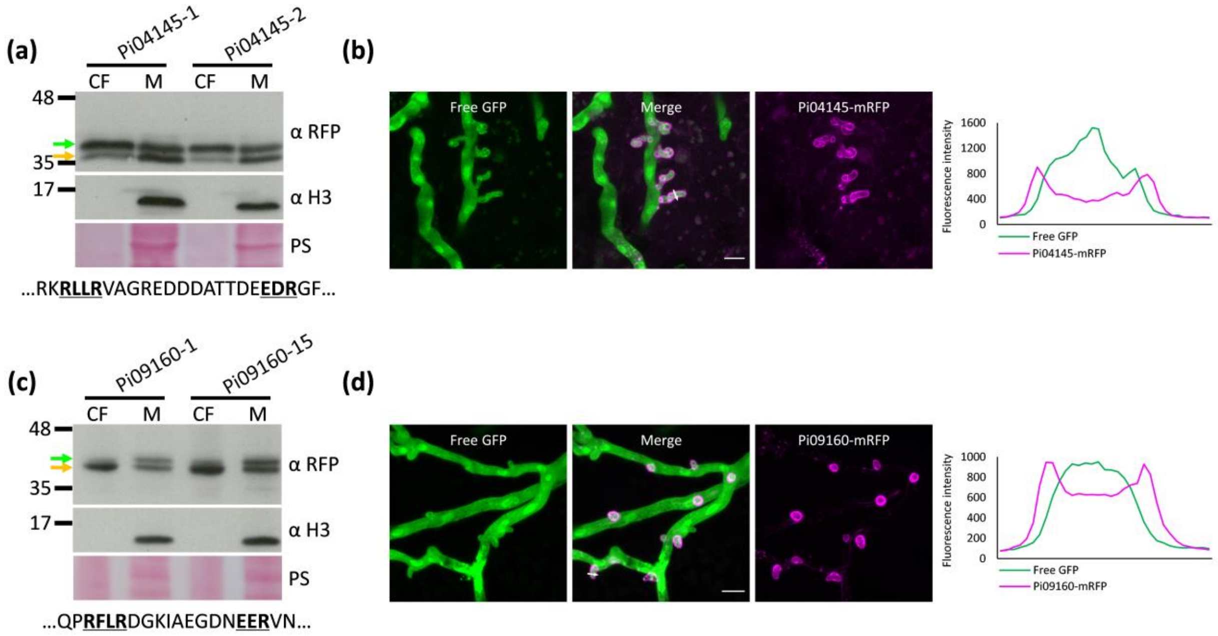


**Figure S9. Secretion of additional RXLR-EER effectors *in vitro* and *in planta*.** Pi04145 (a) and Pi09160 (c) mRFP fusions were detected in the culture filtrate (CF) and mycelium (M) of two different transgenic lines using αRFP. *P. infestans* histone H3 was detected using αH3 only in the M which indicates that the CF was not detectably contaminated by cellular material. Gold arrows indicate fusions cleaved at the EER motif, and green arrows indicate fusions cleaved at the RXLR motif. Markers (kDa) are indicated on the left and protein loading is indicated by Ponceau stain (PS). Partial effector sequences containing the RXLR and EER motifs are shown below the PS. Confocal images of *P. infestans* hyphae and haustoria of Pi04145 (b) and Pi09160 (d) transformants infecting *N. benthamiana* leaves. Effector mRFP fusions were secreted from haustoria and accumulated at the periphery of haustoria (indicated by magenta). Hyphae and haustoria were visualized by the co-expressed cytoplasmic GFP. Images are projections of confocal z-series. Scale bars represent 10 µm. White arrows indicate the lines used for the corresponding fluorescence intensity profiles shown on the right of the images.

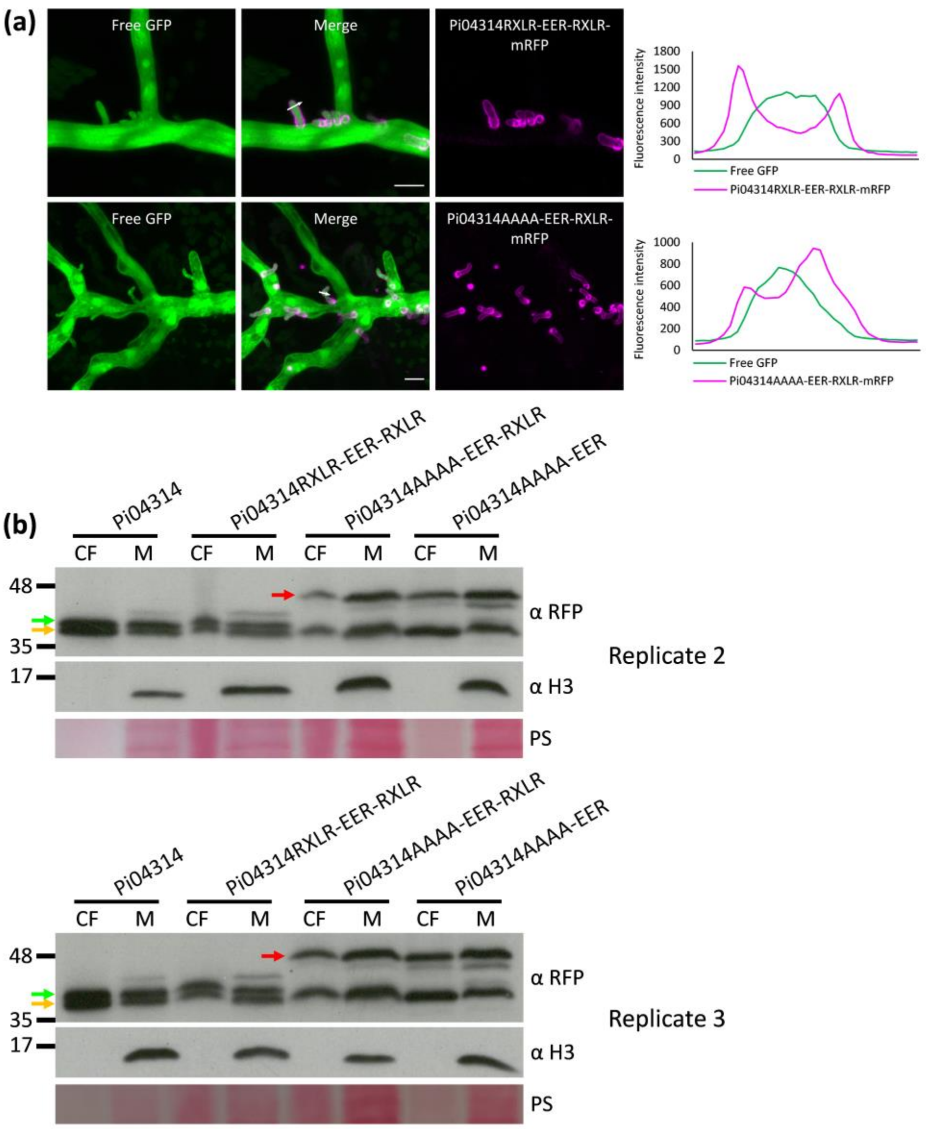


**Figure S10 Secretion of Pi04314RXLR-EER-RXLR and Pi04314AAAA-EER-RXLR *in planta* and *in vitro*.** Confocal images of *P. infestans* hyphae and haustoria of Pi04314RXLR-EER-RXLR (a, upper panel) and Pi04314AAAA-EER-RXLR (a, lower panel) transformants infecting *N. benthamiana* leaves. Pi04314 RXLR-EER-RXLR and AAAA-EER-RXLR mRFP fusions were secreted from haustoria and accumulated at the periphery of haustoria (indicated by magenta). Hyphae and haustoria were visualized by the co-expressed cytoplasmic GFP. Images are projections of confocal z-series. Scale bars represent 10 µm. White arrows indicate the lines used for the corresponding fluorescence intensity profiles shown on the right of the images. (b) Pi04314 WT RXLR-EER-RXLR, AAAA-EER-RXLR and AAAA-EER mRFP fusions were detected in the culture filtrate (CF) and mycelium (M) using αRFP in two independent replicates. *P. infestans* histone H3 was detected using α H3 only in the M which indicates that the CF was not detectably contaminated by cellular material. Gold arrows indicate fusions cleaved at the EER motif, green arrows indicate fusions cleaved at the original RXLR motif, and red arrows indicate fusions cleaved after the SP. Markers (kDa) are indicated on the left and protein loading is indicated by PS.
