## Supplementary Table 1 for "Proteolytic processing of both RXLR and EER motifs in oomycete effectors"

**Supplementary Table S1 Primers used in vector construction for *P. infestans* transformation and *in planta* expression.**

| **Primers used in overexpression vector construction for *P. infestans* transformation** | | |
| --- | --- | --- |
| **Primer name** | **Sequence** | **Used for** |
| 09216-NotI-F | GGAAGCGGCCGCACCATGCGTTTCAGCGTTTTC | PPL-RAG-09216 |
| 09216-NotI-R | GGAAGCGGCCGCGGCGCTGAACCTGTCGTCGT |  |
| 09216RRLR/AAAA-F | AAACGTCGCTGTGGCCAGTGAAGGTGCTGCCGCTGCCGCTGAGCATGCTG | PPL-RAG-09216AAAA |
| 09216RRLR/AAAA-R | CAGCATGCTCAGCGGCAGCGGCAGCACCTTCACTGGCCACAGCGACGTTT |  |
| 04097-up-F | TGCCTCGTCTGACGTTTCTC | Sequence containing Pi04097 |
| 04097-down-R | CCCCGTCACAACGGAGTTTA |  |
| 04097-NotI-F | GGAAGCGGCCGCACCATGCGCAGTATATTCTA | PPL-RAG-04097 |
| 04097-NotI-R | GGAAGCGGCCGCGGCGATGACGATTTCTTAA |  |
| 04097RXLR/AAAA-F | TCAGCTCTTGTCAAAGGCGTCCCCTGACAAAGCAGCCGCTGCGGTCGAAGGCCAAGAA | PPL-RAG-04097AAAA |
| 04097RXLR/AAAA-R | TTCTTGGCCTTCGACCGCAGCGGCTGCTTTGTCAGGGGACGCCTTTGACAAGAGCTGA |  |
| 04314-NotI-F | GGAAGCGGCCGCACCATGCATTCAAGTCTTCTT | PPL-RAG-04314 |
| 04314-NotI-R | GGAAGCGGCCGCGGCGAGTTGGTTTTGTAGAT |  |
| 04314RXLR/AAAA-F | TGAGGAGGAAAACAGCATCGTCGCGGCCG  CCGCCGCAGTCGAGACAAGTGAAGAC | PPL-RAG-04314AAAA-EER |
| 04314RXLR/AAAA-R | GTCTTCACTTGTCTCGACTGCGGCGGCGGCCGCGACGATGCTGTTTTCCTCCTCA |  |
| 04314EER/AAA-F | TCGAGACAAGTGAAGACGAAGCAGCGGCGGATTTGCTTGGGCTTTTTGC | PPL-RAG-04314RXLR-AAA and PPL-RAG-04314AAAA-AAA |
| 04314EER/AAA-R | GCAAAAAGCCCAAGCAAATCCGCCGCTGCTTCGTCTTCACTTGTCTCGA |  |
| 04314KSKL/RSLR-F | AGGGATTTGCTTGGGCTTTTTGCCCGGAGCCTGCGGAAGAAGATGATGAAAAGCGAAAG | PPL-RAG-04314RXLR-EER-RXLR and PPL-RAG-04314AAAA-EER-RXLR |
| 04314KSKL/RSLR-R | CTTTCGCTTTTCATCATCTTCTTCCGCAGGCTCCGGGCAAAAAGCCCAAGCAAATCCCT |  |
| 21388-NotI-F | GGAAGCGGCCGCACCATGCGTTCGCTCCTGTTG | PPL-RAG-21388 |
| 21388-NotI-R | GGAAGCGGCCGCGGGCTAGGGCCAACGTTT |  |
| 21388RXLR/AAAA-F | AACGCCGATGAAAAAGCAGCCGCGGCAGGTGACTACAACAAT | PPL-RAG-21388AAAA-EER |
| 21388RXLR/AAAA-R | ATTGTTGTAGTCACCTGCCGCGGCTGCTTTTTCATCGGCGTT |  |
| 21388EER/AAA-F | CCCAACACGTCTGACGCAGCGGCGGCGTTTTCTATCTCA | PPL-RAG-21388RXLR-AAA and PPL-RAG-21388AAAA-AAA |
| 21388EER/AAA-R | TGAGATAGAAAACGCCGCCGCTGCGTCAGACGTGTTGGG |  |
| 22926-NotI-F | GGAAGCGGCCGCACCATGCTCCGGTCCTTCTTA | PPL-RAG-22926 |
| 22926-NotI-R | GGAAGCGGCCGCGGTGTGGTAAGCTTCGTAAA |  |
| 22926RXLR/AAAA-F | CCCCACAGAAGCACTCATGAAACCGCAGCCGCGGCGACAAACGACGAAGAGAGGGGG | PPL-RAG-22926AAAA-EER |
| 22926RXLR/AAAA-R | CCCCCCTCTCTTCGTCGTTTGTCGCCGCGGCTGCGGTTTCATGAGTGCTTCTGTGGGG |  |
| 22926EER/AAA-F | ATTCCTGAGGACAAACGACGCAGCGGCGGGGGCAACAATGACTTTAG | PPL-RAG-22926RXLR-AAA |
| 22926EER/AAA-R | CTAAAGTCATTGTTGCCCCCGCCGCTGCGTCGTTTGTCCTCAGGAAT |  |
| 22926-7A-F | AGCCGCGGCGACAAACGACGCAGCGGCGGGGGCAACAATGACTTTAG | PPL-RAG-22926AAAA-AAA |
| 22926-7A-R | CTAAAGTCATTGTTGCCCCCGCCGCTGCGTCGTTTGTCGCCGCGGCT |  |
| 04145-NotI-F | GGAAGCGGCCGCACCATGCGCAGTGCATTTTAC | PPL-RAG-04145 |
| 04145-NotI-R | GGAA GCGGCCGCGGATTGCCATCCTTCAGT |  |
| 09160-NotI-F | GGAAGCGGCCGCACCATGCGTCTACCCTCCATC | PPL-RAG-09160 |
| 09160-NotI-R | GGAAGCGGCCGCGGAGCCTTGTTGTTTTGTTC |  |
| mRFP-R | CGCCCTCGATCTCGAACT | For sequencing |
| **Primers used in vector construction for expression *in planta*** | | |
| **Primer name** | **Sequence** | **Used for** |
| SP-09216-gw-F | GGGGACAAGTTTGTACAAAAAAGCAGGCTTCACC ATGCGTTTCAGCGTTTTC | Pi09216_1-175_-mRFP |
| 09216-RXLR-gw-F | GGGGACAAGTTTGTACAAAAAAGCAGGCTTCACC ATGGCTGAGCATGCTGTC | Pi09216_38-175_-mRFP |
| 09216-gw-C-R | GGGGACCACTTTGTACAAGAAAGCTGGGTC CGCTGAACCTGTCGT | Pi09216_1-175_-mRFP and Pi09216_38-175_-mRFP |
| SP-04314-gw-F | GGGGACAAGTTTGTACAAAAAAGCAGGCTTCACC ATGCATTCAAGTCTTCTT | Pi04314_1-154_-mRFP |
| 04314-RXLR-gw-F | GGGGACAAGTTTGTACAAAAAAGCAGGCTTCACC ATGGCAGTCGAGACAAGTGAA | Pi04314_57-154_-mRFP |
| 04314-EER-gw-F | GGGGACAAGTTTGTACAAAAAAGCAGGCTTCACC ATGGATTTGCTTGGGCTTTTTGC | Pi04314_68-154_-mRFP |
| 04314-gw-C-R | GGGGACCACTTTGTACAAGAAAGCTGGGTC CGAGTTGGTTTTGTAGAT | Pi04314_1-154_-mRFP, Pi04314_57-154_-mRFP and Pi04314_68-154_-mRFP |
| SP-21388-gw-F | GGGGACAAGTTTGTACAAAAAAGCAGGCTTCACC ATGCGTTCGCTCCTGTTG | Pi21388_1-152_-mRFP |
| 21388-RXLR-gw-F | GGGGACAAGTTTGTACAAAAAAGCAGGCTTCACC ATGGGTGACTACAACAAT | Pi21388_55-152_-mRFP |
| 21388-EER-gw-F | GGGGACAAGTTTGTACAAAAAAGCAGGCTTCACC ATGGCGTTTTCTATCTCA | Pi21388_73-152_-mRFP |
| 21388-gw-C-R | GGGGACCACTTTGTACAAGAAAGCTGGGTC GCTAGGGCCAACGTT | Pi21388_1-152_-mRFP, Pi21388_55-152_-mRFP and Pi21388_73-152_-mRFP |
| M13-F | GTAAAACGACGGCCAG | For sequencing |
| M13-R | CAGGAAACAGCTATGAC | For sequencing |

NotI restriction enzyme recognition sequence is marked with red, attB1 sequence is marked with blue, attB2 sequence is marked with green, and the start codon is shaded with green.
